# Rho-kinase planar polarisation at tissue boundaries depends on phospho-regulation of membrane residence time

**DOI:** 10.1101/615062

**Authors:** Clara Sidor, Tim J. Stevens, Li Jin, Jérôme Boulanger, Katja Röper

## Abstract

Rho-kinase (Rok) is a major myosin II activator during morphogenesis. In the *Drosophila* embryonic salivary gland placode Rok is planar polarised at the tissue boundary, through a negative regulation by the apical polarity protein Crumbs that is anisotropically localised at the boundary. However, in inner cells of the placode both Crumbs and Rok are isotropically enriched at junctions. We propose a model that reconciles both behaviours through modulation of Rok membrane residence time by Crumbs and downstream effectors. Using FRAP in embryos expressing endogenously-tagged Rok combined with *in silico* simulations, we find that the lower membrane dissociation rate (k_off_) of Rok at the tissue boundary, where Crumbs membrane levels are lower, explains this boundary-specific effect. The S/T-kinase Pak1 negatively affects Rok membrane association *in vivo* within the epidermis, and *in vitro* can phosphorylate Rok near the PH domain that mediates membrane association. Pak1 is recruited to the membrane by Cdc42 which, like its binding partner Crumbs, shows anisotropic localisation at the boundary. These data reveal an important mechanism of modulation of Rok membrane residence time via affecting the k_off_ that may be widely employed during tissue morphogenesis.

## Introduction

Tissues arise during development through specification of primordia that will then initiate morphogenetic movements (Castelli-Gair Hombria, 2016). Many primordia are epithelial in nature and give rise to tubular epithelial organs, such as lung, kidney, intestine in vertebrates or the equivalent organs in invertebrates. How are epithelial primordia physically set aside from the surrounding tissue, apart from the inductive change in transcription factor expression? We know from studies in the *Drosophila* early embryonic epidermis as well as in larval wing discs that differently fated compartments are physically clearly segregated and cell mixing across compartment boundaries is restricted (Dahmann and Basler, 1999; Tepass et al., 2002). In both tissues, this seems to be in part achieved through an increased tension at the compartment boundary that coincides with junctional accumulation of actomyosin into a seemingly supra-cellular structure, a so-called actomyosin cable (Röper, 2013). In the embryonic epidermis, the actomyosin cables found at parasegmental boundaries physically restrains boundary-challenging divisions within the correct compartment (Monier et al., 2010). The molecular mechanisms that drive actomyosin cable assembly are not well understood, and where aspects have been uncovered, a variety of tissue-specific mechanisms seem to contribute. Parasegmental cables for instance arise over time from dorso-ventrally-polarised junctional myosin accumulations (Tetley et al., 2016) that might themselves depend on a code of Toll receptor expression within the early epidermis (Pare et al., 2014). In the wing disc, the dorso-ventral boundary requires Notch-signaling (Major and Irvine, 2005, 2006). Actomyosin-based compartment boundaries are not restricted to invertebrates, but have in fact also been found in vertebrates, with key examples being the rhombomere boundaries in the mammalian hindbrain (Calzolari et al., 2014) as well as the neural plate-ectoderm boundary during neurulation (Galea et al., 2017).

We have previously identified that an actomyosin cable is positioned at the boundary of the salivary gland placode in the *Drosophila* embryo (Röper, 2012). Two epithelial placodes of about 100 cells on either side of the ventral midline become specified at stage 10 of embryogenesis and will invaginate to form the two salivary glands ((Girdler and Röper, 2014; Sidor and Röper, 2016); Fig. 1A,A’). The placodes are contained within parasegment 2 of the embryo, and remnants of previous parasegmental actomyosin cables are specifically retained near the ventral portion of the placode, whilst a new section forms around the dorsal part, so that by mid stage 11 a circumferential actomyosin cable surrounds each placode (Röper, 2012). The formation of this cable is transcriptionally initiated, as it is lacking in mutants for the most upstream specifying transcription factor Sex Combs Reduced (Henderson and Andrew, 2000; Röper, 2012). We have previously identified the apical transmembrane protein Crumbs as a key determinant of actomyosin cable positioning at the boundary of the placode. Crumbs levels are strongly increased within the placode, whereas levels are reduced in the surrounding epidermal cells surrounding the placode. Crumbs’s ability to form homophilic interactions (Fletcher et al., 2012; Röper, 2012; Zou et al., 2012) leads to a highly anisotropic distribution of Crumbs apically within placodal cells at the placode boundary: this Crumbs anisotropy leads to accumulation of actomyosin and also Rho-kinase (Rok) reporters at the placode boundary, away from membranes with high levels of Crumbs. Moreover, ectopic boundaries of high/low Crumbs levels lead to ectopic cable formation (Röper, 2012). Thus, Crumbs exerts a negative regulatory effect on actomyosin cable formation likely through the upstream myosin activator Rho-kinase. Based on studies in mammalian cells that reported that aPKC is able to phosphorylate and thereby inactivate Rok (Ishiuchi and Takeichi, 2011), we previously suggested that the effect in the placode was mediated by aPKC, which in fly embryos closely follows Crumbs distribution (Röper, 2012). Complementary localisation of Crumbs and actomyosin is not restricted to flies, but has also more recently been reported in mouse (Ramkumar et al., 2016).

**Figure 1.**
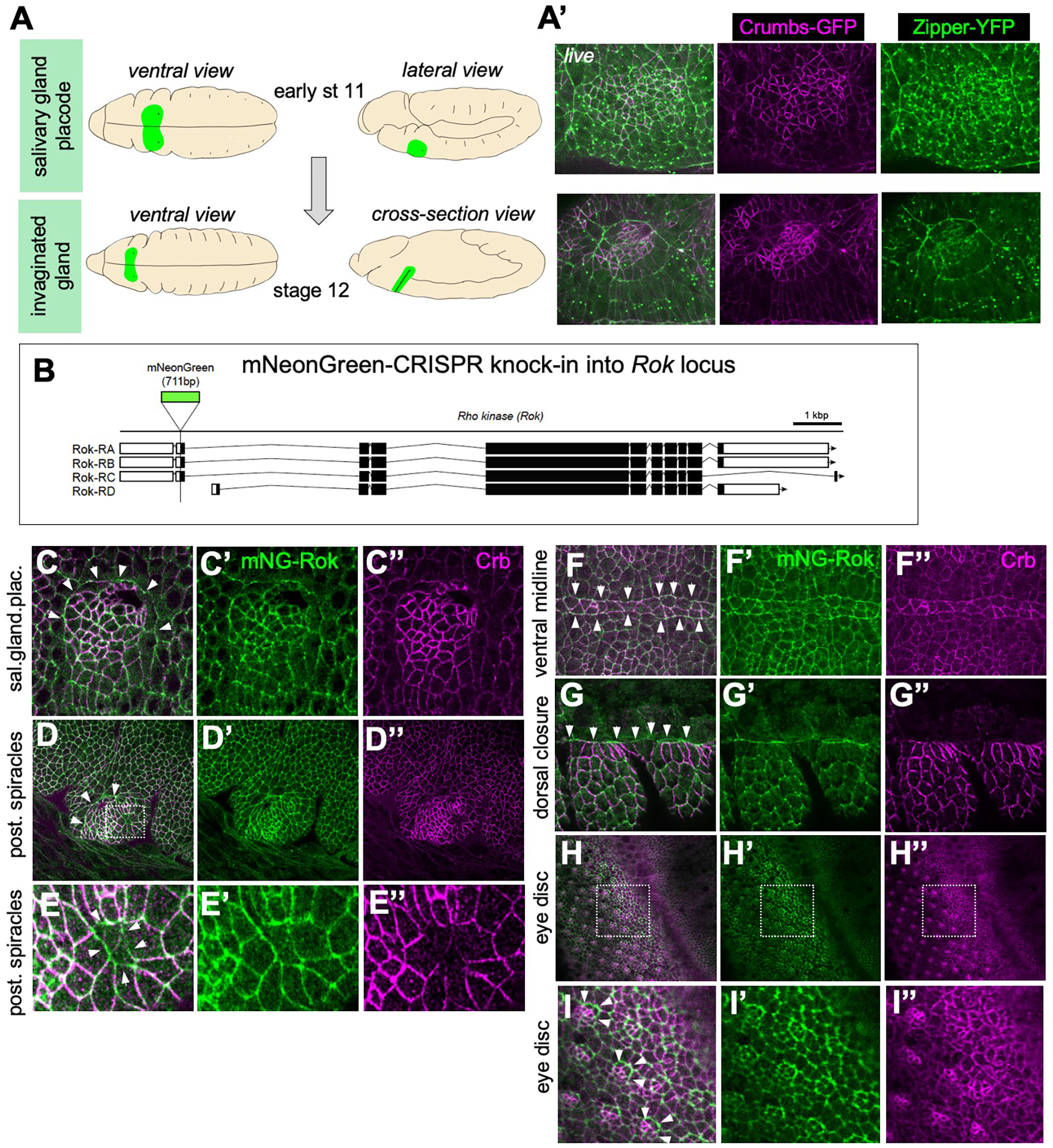
Wide-spread apical planar polarisation of Crumbs and Rho-kinase during morphogenesis. **A** The salivary gland develops from an epithelial placode on the ventral side of the embryo. Cells become specified at late stage 10/early stage 11 (top panels) and by stage 12 most secretory cells have invaginated to form a tube inside the embryo (bottom panels). **A’** The salivary gland placode boundary shows a strong enrichment of myosin II (*Zipper-YFP*, green) into a supracellular cable, complementary to anisotropic enrichment of Crumbs (*Crumbs-GFP*, magenta) in the placodal boundary cells (Röper, 2012); stills of a time lapse movie of matching stages to schematics in **A**. **B** Genomic locus of *Drosophila melanogaster* Rho-kinase (Rok), indicating exon usage in splice variants and the position of the mNeonGreen (mNG) tag inserted at the N-terminus using CRISPR/Cas9. **C-H** mNG-Rok enrichment (green) matching myosin II enrichment and complementary to Crumbs anisotropic localisation (magenta) is found at many epithelial boundaries: the embryonic salivary gland placode (**C-C”**), the embryonic boundary of the posterior spiracle placode (**D-D”**), the embryonic spiracular hair precursors (**E-E’**), the embryonic ventral midline (**F-F”**), the leading edge during embryonic dorsal closure (**G-G”**) as well as the boundaries of specified photoreceptor clusters during larval eye disc morphogenesis (**H-I”**). The boxes in **D, H** indicate magnifications in **E-E”** and **I-I”**, respectivley. Arrowheads point to the boundaries in Crumbs levels that shows strong mNG-Rok accumulation.

The precise role of the actomyosin cable surrounding the salivary gland placode is not clear yet, though we know that it is under increased tension (Röper, 2012). In analogy to the suggested roles of other actomyosin cables, it could actively help tissue invagination by exerting a centripetal force onto the placode tissue. Alternatively, it could actively pull the surrounding epidermis inwards to cover the previous placode area, or it could form an insulating stiff barrier between morphogenetically different domains.

Crumbs is not the only homophilic plasma membrane receptor that can influence actomyosin accumulation near junctions. Depending on the tissue, both E-Cadherin as well as the fly nectin Echinoid show a similar influence at boundaries of expression levels (Chang et al., 2011). Thus, in addition to their adhesive function or, as in the case of Crumbs, their apical polarity role, these receptors have a crucial second role in planar patterning of cytoskeletal structures within the apical domain of epithelial cells.

The negative influence of Crumbs on Rok localisation and activity is clear at the placode boundary where Crumbs anisotropy triggers the actomyosin cable assembly in boundary junctions with lower levels of Crumbs. However, in the inner placodal cells both Crumbs and Rok membrane levels are high and isotropic. High levels of Rok in inner placodal cells drive the tube formation process through a combination of isotropic constriction near the invagination point, mediated by dynamic pools of apical-medial actomyosin, and directed cell intercalation, mediated by junctional pools of actomyosin (Booth et al., 2014; Sanchez-Corrales et al., 2018). Thus, it appears that it is not the overall levels of Crumbs that negatively affect Rok, but rather the difference in levels experienced within a single cell at the boundary that allows negative regulation of Rok. The fact that absolute levels are irrelevant is also supported by the fact that introduction of a new step change in Crumbs levels within the placode, now adding a stripe of Crumbs expression even higher than the already elevated placodal levels, triggers Crumbs anisotropy and ectopic myosin cable formation (Röper, 2012). Here we propose a molecular mechanism by which these different scenarios can be reconciled, based on a modulation of Rok residence time at the apical plasma membrane mediated. We suggest that such mechanism might be widely used in membrane receptors to allow combination of patterning activity with other molecular functions.

## Results

### Rho-kinase is planar polarised and complementary to Crumbs at the placode and other tissue boundaries

We previously proposed that the apical transmembrane protein Crumbs actively influences myosin II accumulation through a negative regulatory effect on Rok, the most common activator of myosin II during morphogenetic processes (Röper, 2012). However, current visualisation of Rok localisation and activity depends on over-expression of tagged active or kinase-dead versions (Simoes Sde et al., 2010). In order to examine endogenous Rok localisation, we used CRISPR/Cas9 to engineer an endogenously N-terminally tagged Rok with the bright mNeon-Green (mNG) fluorescent protein ((Shaner et al., 2013).; Fig. 1B). The tag was inserted at the N-terminal end of the protein in frame with the start, therefore tagging major isoforms A, B and C. Isoform D has an alternative start but it is not expressed during embryogenesis (mod et al., 2010). Several transgenic lines expressing mNG-Rok were obtained. Importantly, mNG-Rok did not form aggregates like many of the overexpression lines (see Fig.4B) and the mNG-Rok flies were homozygous viable, indicating that Rok function was not impaired by the mNG-tag.

In epithelial cells of the epidermis at different stages of embryogenesis as well as in larval imaginal discs, mNG-Rok localised similarly to previously described tagged Rok, being enriched in the apical domain of the cells (Fig. 1C-H; Suppl.Fig.1C). At several boundaries of differently fated epidermal domains, such as the embryonic salivary gland placode (Fig. 1C), the embryonic posterior spiracles (Fig. 1 D, E), embryonic epidermal leading edge/amnioserosa interface (Fig. 1F) and larval eye disc (Fig. 1H, I), mNG-Rok was strongly enriched at the boundary, as was the downstream morphogenetic effector myosin II (Jacinto et al., 2002; Röper, 2012). These were also all boundaries where Crumbs levels show a clear step change of high versus low expression.

### Crumbs induces Rho kinase planar polarisation at the placode boundary

In the embryonic salivary gland placode, mNG-Rok was enriched apically in comparison to the surrounding embryonic epithelium (Fig. 1C and Fig. 2A-A”). However, in placode boundary cells, where Crumbs protein is highly anisotropically localised within the apical domain (Fig. 2A, arrow, and C), Rok was planar polarised (Fig.2A’ arrow). In these boundary cells, mNG-Rok accumulated in junctions with lower levels of Crumbs and appeared depleted from junctions with high Crumbs levels (Fig. 2C), suggesting that Crumbs negatively regulates endogenous Rok membrane accumulation. In order to examine the effect of Crumbs on endogenous Rok membrane localisation, we induced Crumbs overexpression in a stripe of cells in mNG-Rok embryos using the *en-Gal4* driver (using the UAS/Gal4 system; (Brand and Perrimon, 1993)). This ectopic stripe of higher Crumbs levels created a new high/low boundary of Crumbs protein within the salivary gland placode (Fig.2B, arrow). At the boundary of the ectopic stripe, Crumbs localisation was highly anisotropic, with Crumbs enriched in junctions with neighbouring cells that also expressed high levels of Crumbs, and lower at junctions with the surrounding placode cells (Fig. 2 B). Again, only within the cells at the stripe boundary, where Crumbs was highly anisotropic, mNG-Rok was planar polarised and enriched at the new ectopic boundary junctions with lower levels of Crumbs, and it was depleted from junctions with higher levels of Crumbs (Fig. 2 B,’B”). This negative effect was also visible within the apical-basal extent of the epidermal cells, as an expansion of the apico-lateral distribution of Crumbs due to ectopic overexpression lead to a basal shift in mNG-Rok localisation within the lateral membrane (Suppl. Figure 1). Thus, Crumbs negatively regulates endogenous Rok membrane association.

**Figure 2.**
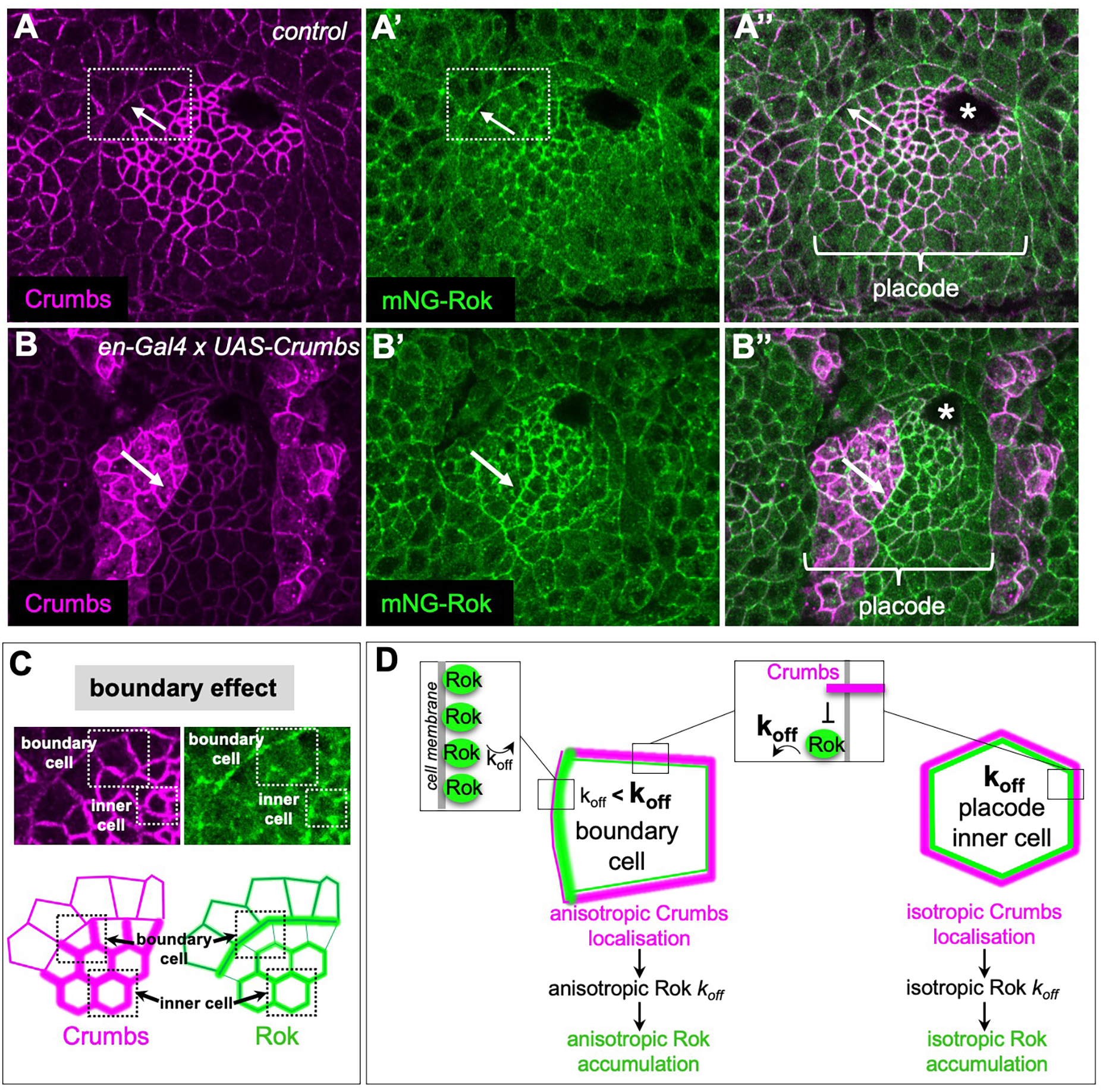
Rho-kinase planar polarisation at the boundary of the salivary gland placode. **A-A”** Crumbs is highly enriched in the salivary gland placode and reduced in levels in the surrounding epidermal cells (**A**, magenta), leading to its strong anisotropic localisation in boundary cells due to homophilic interactions (white arrows in **A-A”**). mNG-Rok is enriched complementary to Crumbs in boundary cells, but also strongly enriched isotropically in junctions in the inner cells of the placode (**A’**, green). **B-B”** Introduction of a new boundary of Crumbs protein levels within the salivary gland placode using *en-Gal4* driven expression of *UAS-Crumbs* (**B**, magenta, levels are adjusted to avoid oversturation in the overexpression stripe) leads to accumulation of mNG-Rok at the new boundary (**B’**, green), complementary to anisotropic Crumbs at the new boundary. See also Supplemental Figure 1. **C** The anisotropic and complementary localisation of Crumbs and mNG-Rok shows a ‘boundary effect’, in that the negative regulatory effect that Crumbs exerts on Rok localisation is only apparent in cells that show Crumbs anisotropy (boundary cell), whereas isotropic Crumbs accumulation in the centre of the placode (inner cells) does not prevent high levels of junctional Rok accumulation. **D** A dynamic model of modulation of Rok residence time at the membrane, depending on local levels of Crumbs, could explain planar polarisation of Rok in cells showing Crumbs anisotropy whilst preserving isotropic Rok localisation in cells with isotropic Crumbs. Asterisks indicate the invagination point.

### A model to explain Rok polarisation at the boundary via modulation of Rok dynamics

We were intrigued by the fact that the negative regulatory effect of Crumbs appeared to only affect Rok at the boundary of the salivary gland placode, but not all throughout the tissue. In inner placodal cells, where Crumbs was highly enriched isotropically at all sub-apical junctions, mNG-Rok was able to localise at these junctions despite the high Crumbs levels (Fig. 2A, C). In contrast, in boundary cells where Crumbs localisation was highly anisotropic, mNG-Rok was planar polarised. Furthermore, an ectopic Crumbs boundary with much higher levels of Crumbs present could also polarise Rok (Fig. 2B-B”). Thus, it was the anisotropic distribution of Crumbs within a cell, rather than its absolute levels in the plasma membrane, that affected Rok membrane localisation (Fig. 2C).

One way to mechanistically reconcile these two situations (boundary versus inner placode) is to consider the dynamics of Rok membrane localisation (Fig. 2D). If we assume that at equilibrium Rok is able to associate and dissociate from the cell membrane at specific rates, Crumbs could influence Rok membrane accumulation by selectively increasing the Rok membrane dissociation rate (k_off_). We propose that within inner placodal cells, high levels of Crumbs in all junctions would lead to a higher, though isotropic, Rok turnover. With Rok membrane recruitment remaining unaffected, Rok would still localise at the plasma membrane in these inner cells. Across the tissue, Rok would dissociate more often from junctions with high levels of Crumbs and would therefore accumulate at junctions with lower levels of Crumbs and lower k_off_, resulting in the planar polarisation of Rok at the tissue boundary.

### Endogenous Rok k_off_ is lower at the boundary of the placode and leads to planar polarisation of Rok in simulations

The above model clearly predicts that Rok’s mobility and in particular Rok’s k_off_ at the placode boundary should differ from Rok’s k_off_ in an inner placode cell. To test this prediction, we performed Fluorescence Recovery After Photobleaching (FRAP) experiments in stage 11-12 embryos expressing the endogenously tagged mNG-Rok (Fig. 3A,B). We bleached small circular region of mNG-Rok at the apico-lateral plasma membrane and imaged at 1s intervals to capture the mNG-Rok fluorescence intensity pre- and post-bleach. mNG-Rok located at junctions within the centre of the placode recovered significantly faster than mNG-Rok located at the boundary (Fig. 3 B-D). k_off_ values were estimated from fluorescence recovery as 0.145 (+/- 0.010) for the inner placodal cells and 0.106 (+/- 0.008) for the boundary and found significantly different using a bootstrap procedure (p-value [bootstrap/boxplot]: 0.00249; Fig. 3C,D). Thus, the dynamic behaviour of mNG-Rok at junctions forming the boundary of the placode compared to inner placode junctions supports the model for Rok planar polarisation presented above.

**Figure 3.**
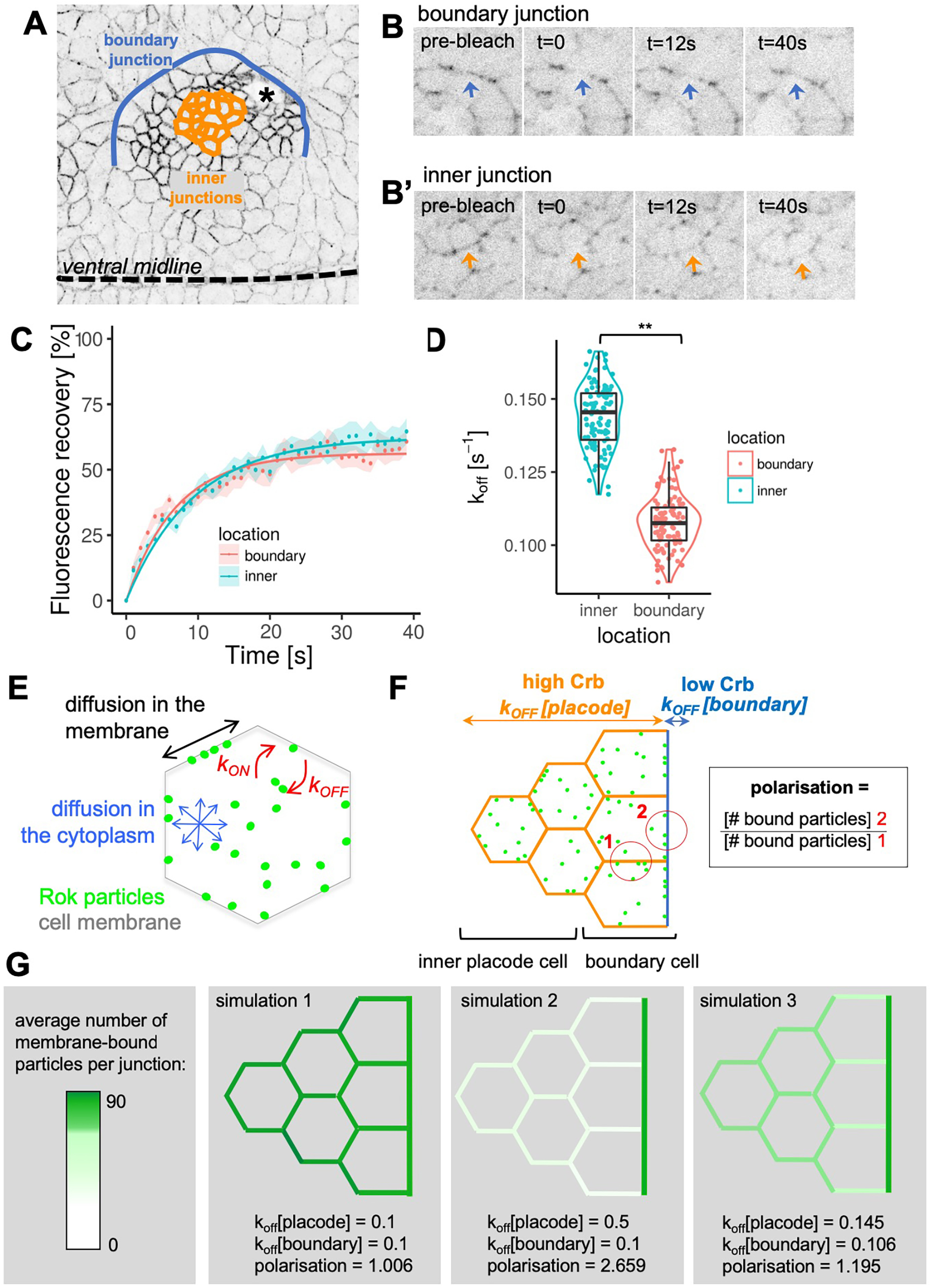
FRAP analysis and in silico simulation of Rok dynamics at the placode boundary versus inside the placode. **A** Schematic of the placode indicating the boundary with low levels of Crumbs (blue) as well as inner placodal cells with isotropically high levels of Crumbs (orange) where FRAP analysis was performed. The invagination point (asterisk) and ventral midline position are indicated. **B, B’** Examples of boundary (**B**) and inner (**B’**) cell junctions of mNG-Rok embryos during FRAP analysis, arrows indicate the positions of the bleached regions (bleach at t=0). **C, D** Recovery curves fitted to data of FRAP experiments for boundary junctions (red; n=26) and inner cell junctions (blue; n=17)(**C**). k_off_ values were estimated from the fluorescence recovery for the boundary as 0.106 (+/- 0.008) and for the inner placodal cells as 0.145 (+/- 0.010) (**C**) and found to be significantly different using a bootstrap procedure (**D**), with the p-value determined as 0.00249 (**). Data represented are bootstrap sample, median and quartiles. **E** We modelled the Rok planar polarisation *in silico* using particle based stochastic reaction-diffusion (see Materials and Methods for details). **F** A group of cells representing boundary and inner placodal cells are modelled, imposing different k_off_ values for the boundary with low Crumbs levels (region 2, blue) and a higher k_off_for membrane with high levels of Crumbs (region 1 for placodal cell, orange). **G** Examples of steady state outputs of the simulation, with simulation 1 depicting no difference in k_off_, simulation 2 assuming a 5-fold difference in k_off_, and simulation 3 using the k_off_ values determined by FRAP as input. See also Supplemental Figure 2.

In order to assess whether the difference in Rok’s k_off_ measured at the placode boundary and within the inner placode is in itself sufficient to produce planar polarisation of Rok at the tissue boundary, we developed an *in silico* simulation of the process, using particle-based stochastic reaction-diffusion (Fig. 3F,G; for details of the implementation see Methods). We tested a variety of k_off_ combinations for the boundary and inner placodal cell membranes (keeping the k_on_ constant; Suppl.Fig. 2). Uniform k_off_ values in all membranes led, as expected, to isotropic membrane accumulation of Rok (Fig. 3G, simulation 1). Setting a high k_off_ in inner junctions compared to boundary junctions led to a strong Rok polarisation at the boundary, but markedly reduced Rok membrane localisation in inner cells (Fig.3G, simulation 2). By contrast, k_off_ values deduced from the *in vivo* FRAP measurements of mNG-Rok were sufficient to trigger Rok planar polarisation in simulated boundary cells, at a level similar to the polarisation measured *in vivo* (*in silico* polarisation value of 1.2, Fig. 3G, simulation 3; compare to values for mNG-Rok *in vivo* in Fig. 6E and Fig. 7G). In these conditions, Rok accumulation at the membrane of inner cells with higher k_off_ was preserved, leading to a pattern that closely resembled Rok localisation in the salivary gland placode *in vivo* (Fig. 1C’ and 2 A’).

Thus, modulation of Rok membrane k_off_ within the placode is sufficient to elicit Rok planar polarisation specifically at the tissue boundary without affecting Rok accumulation in the rest of the tissue. We next investigated the molecular basis for this mechanism.

### The Rok C-terminal region is required for its planar polarisation

*Drosophila* Rok is a large protein of 1391aa, containing an N-terminal kinase domain followed by a coiled-coil region, and a C-terminal Shroom-binding domain (SBD), Rho-binding domain (RBD) and PH-domain (PH) (Fig. 4A)(Amano et al., 2010; Simoes Sde et al., 2014). In order to identify which of these domains were required for apical planar polarisation, truncated versions of a Venus-tagged kinase-dead Rok localisation reporter were overexpressed in embryos using the UAS/Gal4 system under the control of *Daughterless-Gal4*, an early ubiquitous zygotic driver (Simoes Sde et al., 2014)

**Figure 4.**
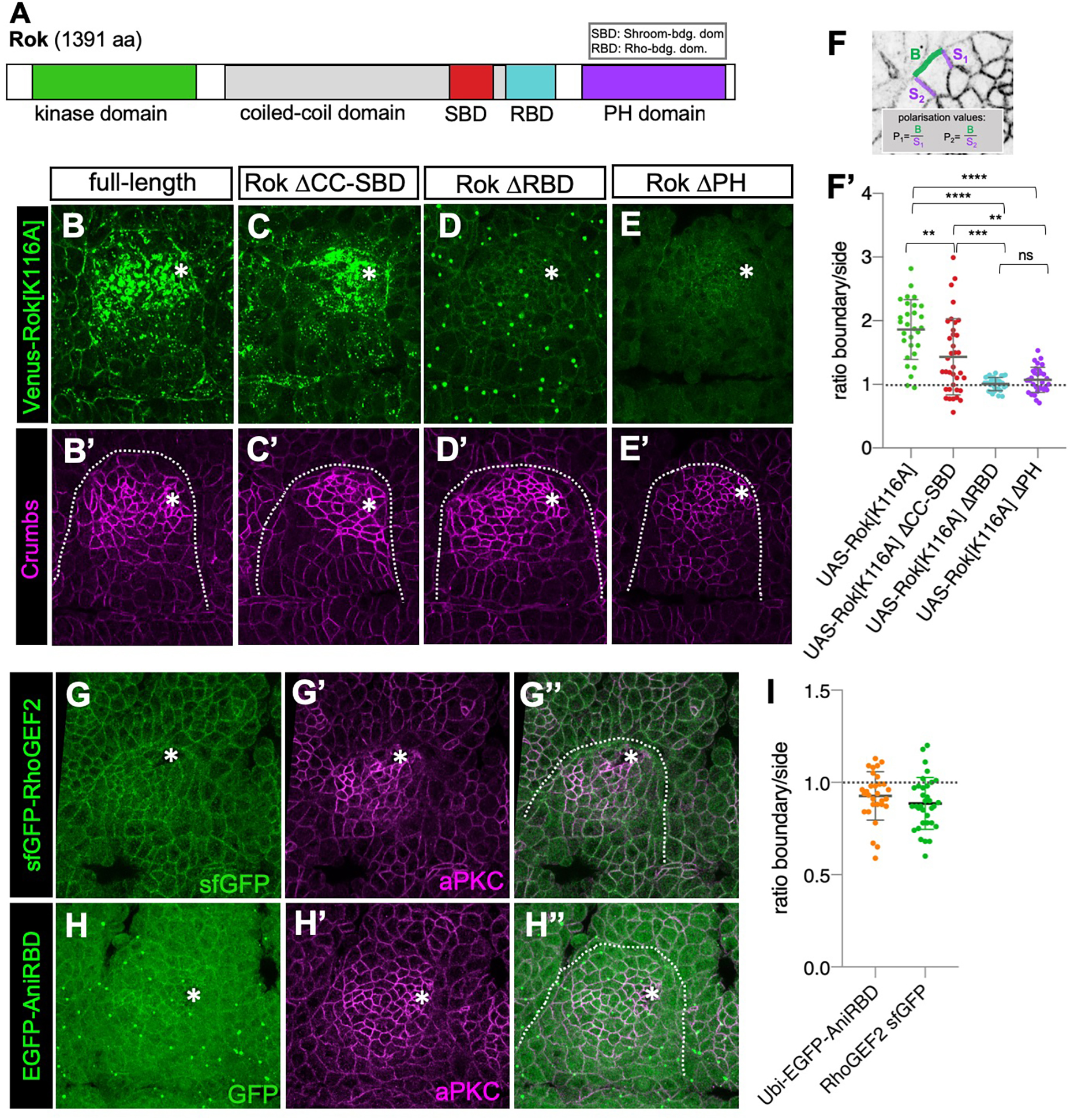
Rok’s RBD and PH domain are required for its planar polarisation and placode enrichment. **A** Schematic of Rok protein domains: N-terminal kinase domain followed by a coiled-coil domain, a Shroom-binding domain (SBD), a Rho-binding domain (RBD) and a C-terminal PH-domain. **B-F’** Expression of variations of a Venus-tagged and kinase-dead localisation reporter of Rok (*Venus-Rok[K116A]*) throughout the embryo using *Da-Gal4*: **B, B’** full-length Venus-Rok[K116A] looks similar to endogenously tagged mNG-Rok and shows anisotropic enrichment at the boundary (**F’**), though note that its accumulation at certain sites such as in the apical-medial region is enhanced, likely due to the overexpression. **C, C’** Rok lacking the coiled coil and SBD (Rok ΔCC-SBD) is less polarised at the boundary but retains some anisotropy (**F’**) and also shows some apical-medial aggregation. **D,D’** Rok lacking the RBD (Rok ΔRBD) is still localised to cell junctions but not enriched in the placode and shows no polarisation at the boundary (**F’**). **E,E’** Rok lacking the PH domain (Rok ΔPH) is still localised to cell junctions but not enriched in the placode and shows no polarisation at the boundary (**F’**). **F,F’** Polarisation quantification expressed as the intensity ratio of boundary junction versus side junctions (**F**). **F’** Mean values are: Rok[K116A]=1.86; Rok ΔCC-SBD=1.433; Rok ΔRBD=1.005; Rok ΔPH=1.701. Data are represented as data points, mean and SEM. Statistical tests used were unpaired t-test, p values are indicated with ** being <0.005, *** being <0.0005, ****<0.0001, ns being not significant. **G-I** Neither RhoGEF2 (**G-G”**; visualised using *sfGFP-RhoGEF2*) nor active Rho (**H-H”**; visualised using *EGFP-AniRBD*) are polarised at the placode boundary (**I**). **I** Polarisation quantification expressed as the intensity ratio of boundary junction versus side junction. Data are represented as data points, mean and SEM. Mean values are: sfGFP-RhoGEF2=0.9267; EGFP-AniRBD=0.8862.

The Venus-Rok[K116A]-ΔSBD lacking the coiled-coil/Shroom-binding domain was able to accumulate in apical membranes (at lower levels than the control Venus-Rok[K116A]) and could still polarise at the placode boundary (Fig. 4B vs C, and F’). Venus-Rok constructs lacking the Rho-binding domain, Venus-Rok[K116A]ΔRBD, or the PH-domain, Venus-Rok[K116A]ΔPH, although showing a strongly reduced overall membrane accumulation, were able to localise at low levels to the membrane. Both constructs failed to polarise at the tissue boundary (Fig. 4D-E’, and F’). Thus, in addition to promoting Rok apical membrane recruitment, both the Rho-binding domain and the PH domain are important for Rok planar polarisation in boundary cells.

Rok membrane recruitment has been shown to be dependent on the small GTPase Rho and its exchange factor RhoGEF2 (Amano et al., 2010; Mason et al., 2016; Nakamura et al., 2017). In order to assess whether Rok planar polarisation is caused by planar polarisation of these upstream activators, we examined the localisation of an sfGFP tagged version of RhoGEF2 (Sarov et al., 2016) and the GFP tagged Rho-binding domain of Anillin (AniRBD-GFP), a reporter for activated GTP-bound Rho (Munjal et al., 2015). Neither RhoGEF2-sfGFP nor AniRBD-GFP showed any apical planar polarisation within the boundary cells of the salivary gland placode (Fig. 4 G-I). Thus, Rok planar polarisation is not driven by an upstream polarisation of active Rho.

Taken together, these data suggest that, although Rho-binding was important for Rok membrane recruitment, once at the membrane Rok planar polarisation is regulated by a Rho-independent mechanism involving the Rok C-terminal membrane association region.

### Phosphorylation of the Rok C-terminal region by Pak1 regulates Rok membrane association

Alternatively, regulation of Rok membrane association could be achieved via modulation of Rok’s binding to Rho or to phospholipids, for instance through phosphorylation in the vicinity of Rok’s RBD or PH domain. Phosphorylation near phospholipid-interacting sequences in several Par-complex substrates has recently been suggested to inhibit their membrane binding (Bailey and Prehoda, 2015). We had previously suggested that Crumbs could negatively regulate Rok through one of its downstream interactors, aPKC (Fig. 5A; (Röper, 2012)), which binds to the Crumbs intracellular domain through its binding partner Par6 (Bulgakova and Knust, 2009). Previous data from mammalian tissue culture cells revealed that in this *in vitro* system aPKC could phosphorylate human ROCK1, one of the two mammalian Rho-kinases, near the RBD and PH-domains and a phospho-mimetic version of ROCK1 showed loss of plasma membrane association (Ishiuchi and Takeichi, 2011). Furthermore, the p21-activated kinase 1 (Pak1), which is activated by the Par6 binding protein Cdc42, has recently been shown to act semi-redundantly with aPKC to phosphorylate shared target proteins (Fig. 5A; (Aguilar-Aragon et al., 2018; Bokoch, 2003)). Importantly, several putative phosphorylation sites for both Pak1 and aPKC are highly conserved between *Drosophila* and mammalian Rho kinases (Blom et al., 1999; Blom et al., 2004; Rennefahrt et al., 2007); Fig. 5B, Suppl. Fig.3). We therefore set out to examine whether aPKC and/or Pak1 contributed to a regulation of Rok membrane association.

**Figure 5.**
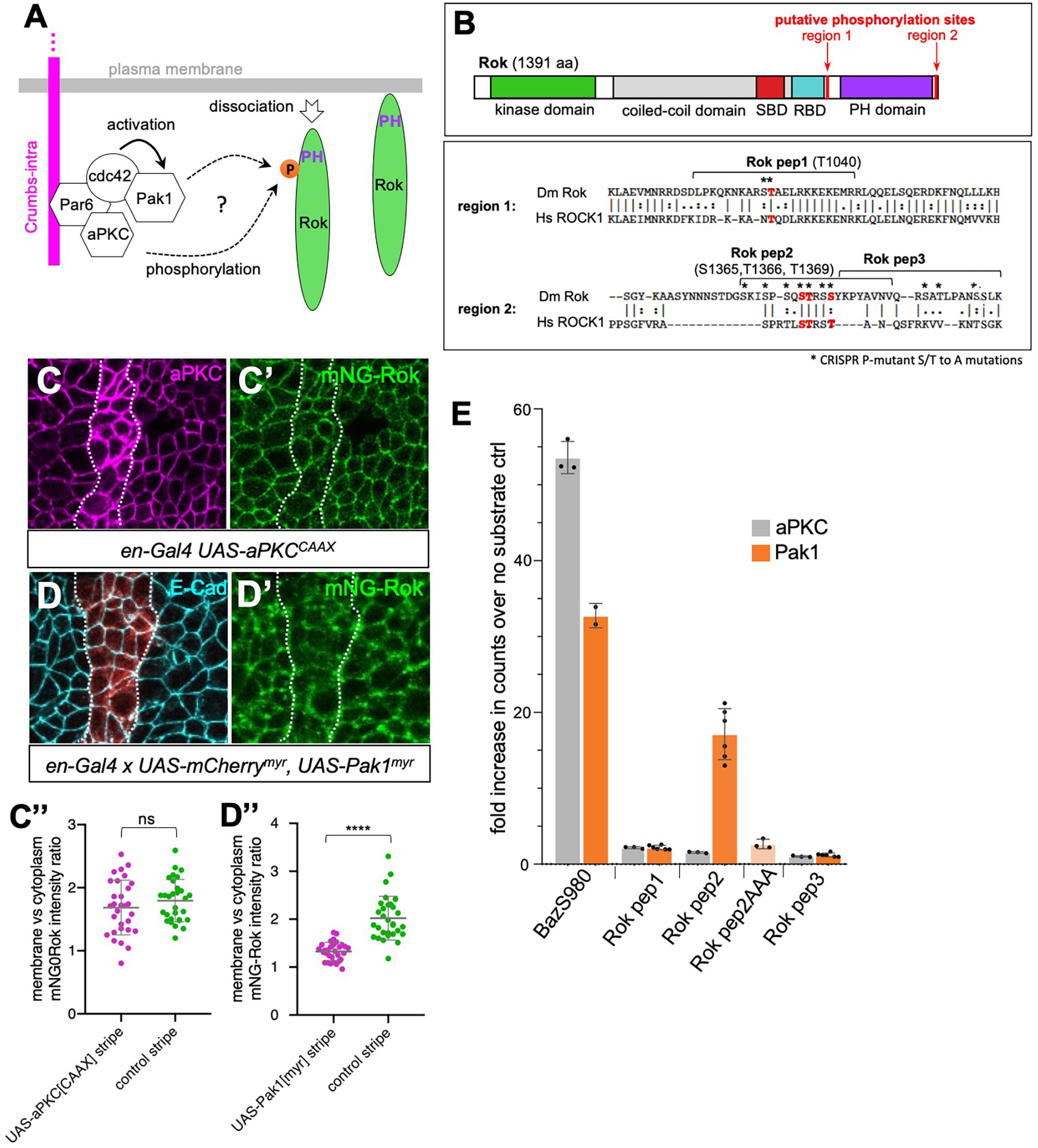
Pak1 can phosphorylate Rok and induce its dissociation from the membrane. **A** Crumbs’ intracellular tail can interact with two kinases with highly overlapping phosphorylation targets: aPKC and Pak1. **B** Both aPKC and Pak1 have many overlapping predicted phosphorylation sites in Rok, in particular near the C-terminal RBD and PH domains. Residues marked in red and named above the sequence are conserved between *Dm* Rok and *Hs* ROCK1 and are phosphorylated in human EpH4 cells (Ishiuchi and Takeichi, 2011). Residues marked by asterisks were mutated in mNG-Rok[Pmut], see Fig.7. We propose that phosphorylation of Rok by either or both kinases could promote its dissociation from the plasma membrane. See also Supplemental Figure 3. **C-C’** Overexpression in stripes of a membrane-targeted version of aPKC (magenta in **C**; using *en-Gal4 UAS-aPKC[CAAX]*) does not affect membrane localisation of mNG-Rok (green) in the overexpressing stripes, as quantified in **C”**. Data are represented as data points, mean and SEM. Unpaired t-test was used to determine statistical significance, p value is 0.278 (ns). **D-D”** Overexpression in stripes of a membrane-targeted version of Pak1 (red marks overexpression stripe in **D**; using *en-Gal4 × UAS-mCherry, UAS-Pak1[myr]*) leads to loss of mNG-Rok (green) from the membrane in the overexpressing stripes, as quantified in **D”**. Data are represented as data points, mean and SEM. Unpaired t-test was used to determine statistical significance, p value is <0.0001 (****). **E** Using purified kinase domains of either aPKC or Pak1 and short peptide substrates of Rok (indicated in **B**) in an *in vitro* kinase assay, we can detect phosphorylation of Rok peptide 2 (Rok pep2), that is located near the PH domain, by Pak1, but no phosphorylation of any Rok peptide tested by aPKC. Both kinase domains phosphorylate a known substrate peptide from Bazooka (BazS980). Data show fold enrichment of radioactive P[33]-phosphate counts over ‘no substrate’ control, data points, mean and SEM are shown. Rok pep2AAA is identical to the Rok pep2 peptide, with three potential phosphorylation sites mutated to alanine, S1365A, T1366A, T1369A.

To examine the role of aPKC and Pak1 in Rok membrane localisation, we used the GAL4/UAS system to overexpress membrane targeted versions of aPKC or Pak1 in engrailed stripes in embryos with endogenously tagged mNG-Rok. While overexpression of UAS-aPKC[CAAX] did not significantly affect mNG-Rok membrane localisation in the embryonic epithelium in comparison to control cells (Fig. 5C-C”), overexpression of UAS-Pak1[myr] strongly decreased mNG-Rok membrane localisation (Fig. 5D-D”).

Mass Spectrometric analysis of mammalian EpH4 tissue culture cell lysates performed by (Ishiuchi and Takeichi, 2011) identified nine phosphorylated sites in human ROCK1 in these cells, four of which are conserved in *Drosophila* Rok. Strikingly, all four sites are located in the Rok C-terminal region, close to the RBD and PH domains (Fig. 5B and Suppl. Fig.3). Furthermore, all are recognised as putative phosphorylation substrates for Pak1 and aPKC (Fig. 5B; Suppl Fig. 3). We designed three short peptides covering these putative sites and performed *in vitro* kinase assays with the purified kinase domains of human Pak1 and human PKCι, using a small Bazooka peptide (BazS980) as a positive control (Fig. 5E). Both Pak1 and aPKC kinase domains strongly phosphorylated the control Bazooka peptide, but no aPKC phosphorylation of any of the Rok peptides was detected. By contrast, Pak1 strongly phosphorylated Rok peptide 2 (Rok Pep2), containing serine S1365 and threonines T1366 and T1369, which are all located close to the C-terminal end of the PH domain (Fig. 5B, E). This phosphorylation was completely abolished in a peptide that had these three residues, S1365, T1366 and T1369, mutated to alanine (Fig. 5E; Rok pep2AAA).

Altogether these data suggest that Pak1 negatively regulates Rok membrane accumulation by phosphorylating its C-terminal region.

### Pak1 regulates Rok planar polarisation downstream of Crb/Cdc42

Although zygotic loss of Pak1 is lethal, maternal contribution allows embryos to develop normally as far as early stage 11, making it possible to investigate Pak1’s role during salivary gland placode morphogenesis. Interestingly, Pak1 zygotic loss of function has been shown to induce defects in embryonic dorsal closure as well as in the late embryonic salivary glands (Bahri et al., 2010; Conder et al., 2007; Pirraglia et al., 2010)). In early stage 11 *pak1^14^* zygotic mutant embryos, Pak1 was still detectable in the epithelium and early salivary placodes were not affected. However, in later placodes at stage 12 mNG-Rok planar polarisation at the salivary gland boundary was strongly reduced and the overall organisation of the placode was affected (Fig. 6A-B’). Moreover, UAS-Pak1[myr] overexpression in the salivary placode using fkh-Gal4 (Henderson and Andrew, 2000) completely abolished mNG-Rok planar polarisation at the boundary (Fig. 6C and E) and also induced strong defects at the tissue boundary, most obvious at later stages when the placode seemed to pull away from the surrounding embryonic epithelium (Fig. 6D). These results show that Pak1 plays a crucial role in salivary gland morphogenesis by regulating Rok membrane localisation and planar polarisation.

**Figure 6.**
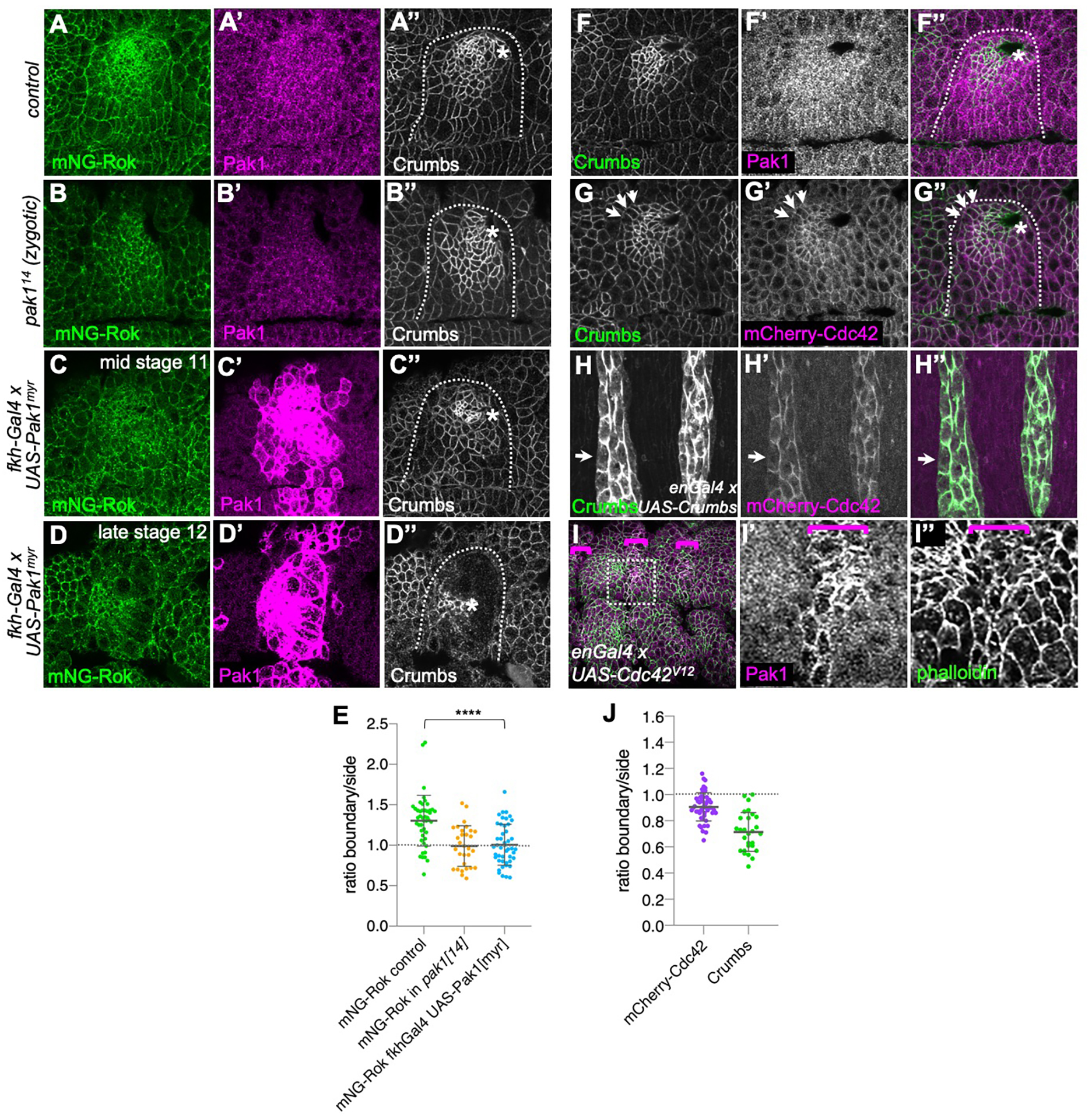
Pak1 acts downstream of Crumbs/Cdc42 and controls Rok planar polarisation and cable formation at the placode boundary. **A-B”** In contrast to the wild-type (**A-A”**) where Pak1 is enriched in the placode (**A’**) and mNG-Rok strongly accumulates at the placode boundary (**A**), in *pak1^14^* zygotic mutants Pak1 antibody labelling is strongly reduced (**B’**) and mNG-Rok is not polarised to the placode boundary (**B**, quantification in **E**). **C-D”** Overexpression of the membrane-targeted form of Pak1, using *UAS-Pak1[myr]*, in the salivary gland placode alone using *fkh-Gal4* leads to a disorganised placode at stage 11 (**C”**) and loss of mNG-Rok polarisation (**C**, and quantification in **E**). At late stage12, many *UAS-Pak1[myr] × fkhGal4* placodes show a disruption that suggests a ripping of cells at the placode boundary (**D”**). mNG-Rok remains at remnant cell boundaries (**D**). **C’** and **D’** show the increased levels of overexpressed membrane-localised Pak1 by antibody staining. **E** Polarisation quantification expressed as the intensity ratio of boundary junction versus side junction. Data are represented as data points, mean and SEM. Mean values are: mNG-Rok control=1.304; mNG-Rok in *pak1^14^*=0.99; mNG-Rok fkh-Gal4 UAS-Pak1[myr]=1.006. Statistical tests used were unpaired t-test, p values are indicated with **** being <0.0001. Dotted lines indicate the placode boundary and asterisks the invagination point. **F-F”** Pak1 (**F’** and magenta in **F”**; revealed by anti-Pak1 antibody) is enriched in the salivary gland placode compared to the surrounding epidermis. Crumbs is shown in **F** and green in **F”**. **G-G”** Cdc42-mCherry localisation (**G’** and magenta in **G”**) follows endogenous Crumbs anisotropic localisation (**G** and green in **G”**) at the placode boundary (arrows). Polarisation quantification of mCherry-Cdc42 and Crumbs in corresponding cells expressed as the intensity ratio of boundary junction versus side junction (**J**). Mean values are: mCherry-Cdc42: 0.906; Crumbs: 0.714. Data are represented as data points, mean and SEM. **H-H”** Overexpression of Crumbs (**H** and green in **H”**) in *enGal4* stripes leads to ectopic recruitment of mCherry-Cdc42 (**H’** and magenta in **H”**) to sites of ectopic Crumbs, again following the anisotropy (arrows). **I-I”** Overexpression of a constitutively active form of Cdc42, Cdc42^V12^, in *enGal4* stripes leads to a strongly increased membrane association of Pak1 (magenta in **I** and single channel in **I’**) in the overexpressing stripes. Membranes are labelled with phalloidin to reveal F-actin (green in **I** and single channel in **I”**). **I’** and **I”** are magnifications of the box indicated in **J**. Dotted lines indicate the boundary of the placode and asterisks mark the invagination point.

To understand how Crumbs might affect Pak1 to control Rok planar polarisation, we examined Pak1 protein localisation in the embryonic epidermis. Like Crumbs protein, Pak1 was localised apically and was enriched in the salivary placode (Fig. 6F-F’). Pak1 function depends on its activation by the small GTPase Cdc42 (Bokoch, 2003). An mCherry-Cdc42 reporter expressed under the control of the *sqh* promoter localised apically in the embryonic epithelium and showed anisotropic distribution at the salivary gland placode boundary similar to Crumbs (Fig. 6G-G”). Ectopic Crumbs expressed in *en-Gal4* stripes also recruited this Cdc42-reporter to ectopic locations (Fig. 6H-H”). In order to confirm that Cdc42 activated Pak1 in the embryonic epidermis we examined the localisation of Pak1 in cells expressing constitutively active Cdc42^V12^ (Welch et al., 1998). In stripes of cells expressing Cdc42^V12^ Pak1 localisation to junctions was strongly enhanced (Fig.6I-I”).

Thus, the negative regulatory effect of Crumbs on Rok is likely mediated by Pak1, recruited and activated by Crumbs-bound Cdc42.

### Phosphorylation of Rok contributes to its planar polarisation in vivo

To confirm that phosphorylation of Rok by Pak1 or aPKC played a role in its planar polarisation *in vivo*, we used the previously generated mNG-Rok strain to mutate the four conserved putative phosphorylation sites described above, as well as eight serines and threonines in close proximity (Fig. 5B, asterisks) using CRISPR/Cas9 and homologous recombination repair (Fig. 7A).

**Figure 7.**
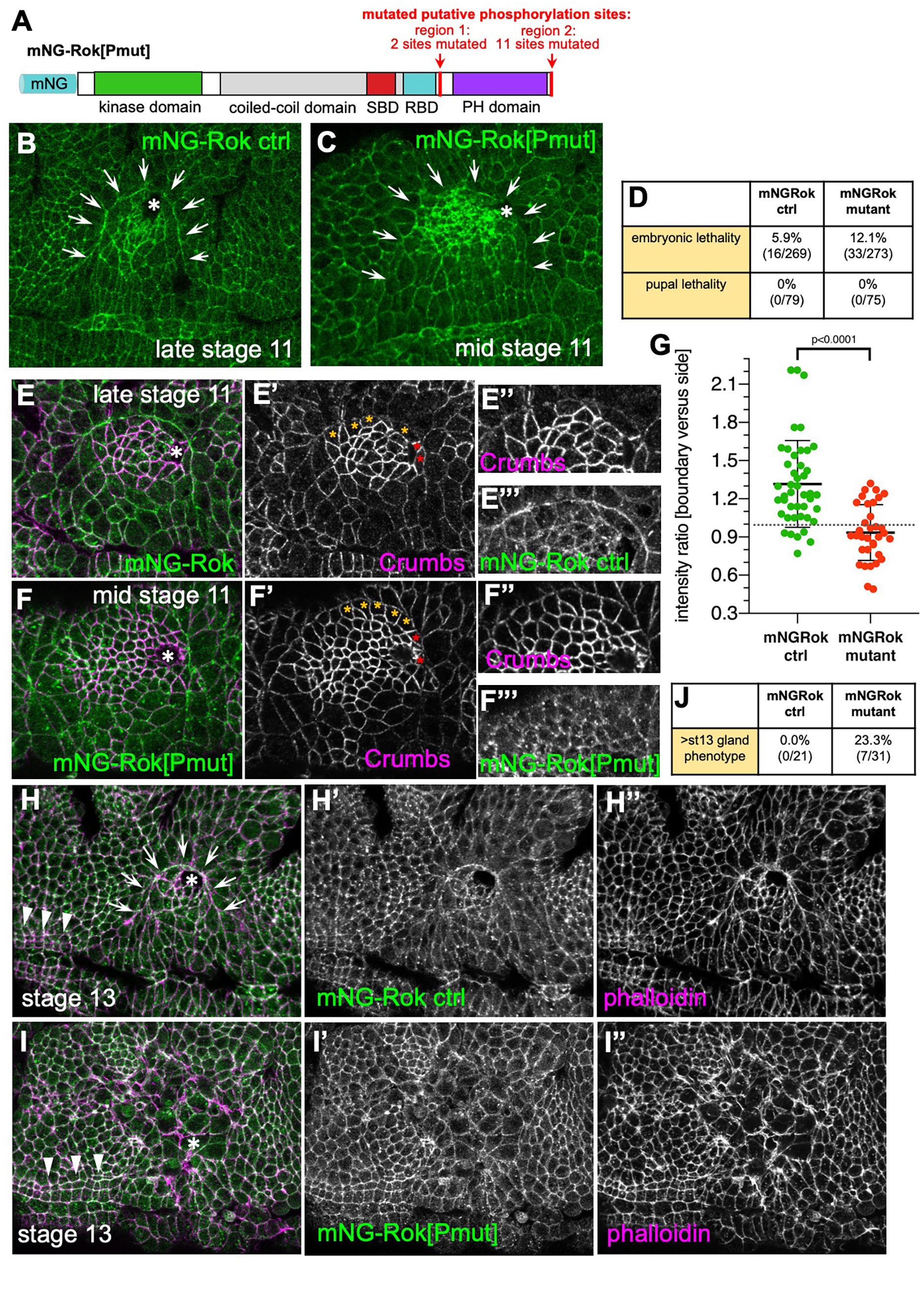
A phospho-mutant Rok shows reduced planar polarisation at the boundary. **A** Schematic of the potential target sites for aPKC or Pak1 phosphorylation that are mutated in the phospho-mutant Rok. **B,C** Localisation comparison between mNG-Rok (**B**) and phospho-mutant Rok (mNG-Rok[Pmut], **C**). Arrows point to the position of the boundary, the asterisks mark the invagination point. **D** Flies homozygous for phospho-mutant Rok are semi-viable, with 12.1% of embryos not hatching compared to 5.9% in the corresponding mNG-Rok control. **E-G** Compared to the mNG-Rok control (green in **E**, and **E”’**) that shows polarisation at the boundary marked by Crumbs (magenta in **E**, and **E’**, **E”**), mNG-Rok[Pmut] (green in **F**, and **F”’**) shows a loss of polarisation at the boundary marked by Crumbs (magenta in **F**, and **F’**, **F”**). **G** Polarisation quantification expressed as the intensity ratio of boundary junction versus side junction. Data are represented as data points, mean and SEM. Mean values are: mNG-Rok ctrl: 1.317; mNG-Rok[Pmut]: 0.935. Unpaired t-test was used to compare samples, p value is indicated. **H-J** At stage 13 phospho-mutant Rok embryos show a strong disruption of the placodal boundary in 23.3% of embryos (compared to 0.0% in the control), Rok is in green in **H**, **I** and single in **H’**, **I’**, with phalloidin to label cell outlines in magenta in **H**, **I** and single in **H”**, **I”**. **J** Quantification of stage 13 gland phenotype prevalence. Arrowheads in **H**, **I** point to the pharyngeal ridges and identify embryos as stage 13, arrows in **H** point to the planar polarised mNG-Rok control at the boundary.

Although flies carrying this mNG-tagged phospho-site-mutant Rok (mNG-Rok[Pmut]; Fig. 7 B vs C) were homozygous viable, a two-fold increase in embryonic lethality (12.1% of fertilised embryos) was observed in comparison to the parental mNG-Rok strain (5.9% of fertilised embryos) (Fig. 7D).There was no increased lethality at later developmental stages. We analysed planar polarisation of mNG-Rok[Pmut] in comparison to mNG-Rok by quantifying fluorescence intensity ratio between boundary/outside membranes and boundary/boundary membranes in cells that showed a clear Crumbs anisotropy (Fig. 7 E-G). Whereas, in agreement with quantifications shown in Fig. 6E, mNG-Rok showed a clear polarisation at the boundary, with a mean ratio of 1.317, mNG-Rok[Pmut] was on average not polarised, with a mean ratio of 0.935 (Fig. 7G). This suggests that phosphorylation of these consensus aPKC/Pak1 sites contributes to Rok planar polarisation.

We could not detect any phenotype in tissue bending and salivary gland placode invagination at early stages, i.e. stages 11-12, in the phospho-mutant Rok embryos, but could detect a fraction of embryos beyond stage 13 (23.3% in mNG-Rok[Pmut] compared to 0.0% in the mNG-Rok control) that showed a striking phenotype: we observed a disruption and altered cell shapes within the epidermis at the positions where the salivary gland placodes were located (Fig. 7H-J).The rest of the epidermis appeared largely unaffected. At this stage in control embryos, the circumferential actomyosin cable at the placode boundary was strongly highlighted when stained with phalloidin to label F-actin (Fig. 7 H,H”; (Röper, 2012)). By contrast, we did not detect any accumulation of F-actin at comparable junctions in the mNG-Rok[P[mut] where the cable would be positioned in the wild-type (Fig. 7 I,I”). Matching the phalloidin distribution, the mNG-Rok[Pmut] was still localised to junctions in the placode area but there was no accumulation suggesting cable localisation at this stage (Fig. 7 I’).

Thus, in agreement with the loss of Rok planar polarisation in the *pak1^14^* mutant, blocking phosphorylation of the Rok C-terminal domain by Pak1 through mutation of the consensus sites affected Rok’s ability to be planar polarised.

## Discussion

The prepatterning of tissues to prime and prepare them for subsequent morphogenetic changes in development is key to successful organ morphogenesis. This is initially controlled at the transcriptional level through morphogenetic networks of transcription factors that drive primordium specification. How such patterning is then transformed into the specific activation and placement of morphogenetic effector proteins that impinge on cell shape and cell behaviour is much less understood.

In epithelial cells, junctional proteins as well as morphogenetically active pools of actomyosin are concentrated within the apical and apico-lateral region of the cells. This placement is controlled by the epithelial polarity network (Tepass, 2012). Patterning of cytoskeletal activity and junctional changes, key ingredients of morphogenetic changes in epithelial tissues, then takes place within this apical domain. This can lead to apical planar polarisation of whole tissues or smaller domains or rows of cells that are morphogenetically active, as for instance is the case at the boundary of the salivary gland placode. In this case, the apical polarity determinant Crumbs plays a dual role, firstly in maintaining apical-basal polarity of epidermal cells including the salivary gland placode, and secondly in patterning cytoskeletal behaviour within the apical domain. Crumbs levels, in fact, show dynamic variations across much of the *Drosophila* embryonic epidermis until stage 14, and as described here, step changes in Crumbs levels tend to be accompanied by actomyosin accumulation at these boundaries (Röper, 2013).

Rok as the key morphogenetic activator of non-muscle myosin II is crucial to development in many animals. Thus, regulation and activity of Rok in cells is closely controlled. Historic views of Rok activation assumed a potential fold-back mechanism whereby the known regulatory activity of Rok’s C-terminus would contact the N-terminal kinase domain and block its activity (Amano et al., 1999; Julian and Olson, 2014). An alternative view is supported by recent evidence and suggests that the C-terminal domain is crucial for membrane interaction and that Rok is in fact always found as a homodimer in an extended conformation (Truebestein et al., 2015; Truebestein et al., 2016). Such an extended conformation and role of the C-terminus in membrane binding is in agreement with our *in vivo* findings that phospho-regulation of this domain is critical for membrane localisation.

aPKC and Pak1 are both important kinases with a multitude of roles in development and tissue homeostasis (Hong, 2018; Rane and Minden, 2014). Their overlapping function though has only been appreciated recently (Aguilar-Aragon et al., 2018). We originally favoured a role for aPKC in mediating the Crumbs regulatory effect on Rok at the salivary gland placode boundary, in particular as aPKC localisation closely mimics Crumbs localisation in the fly embryonic epidermis. Without the ability to visualise endogenous Rok, any previous interpretation of effects of aPKC on Rok were indirect, and overexpression constructs of Rok appear to not completely copy endogenous Rok behaviour. But why ‘charge’ the Crumbs intracellular domain with two kinases with highly overlapping targets? The fact that Pak1 depends on Cdc42 for its activation might add another layer of control and allow differential kinase usage or amplification of kinase activity depending on the tissue context.

The differential expression of Crumbs between salivary gland placode and surrounding epidermis, or epidermis and amnioserosa, demarcates clear boundaries. These boundaries at the plasma membrane level can then be turned into cytoskeletal planar polarisation, leading to physical boundaries due to for instance increased tension at these boundaries. Crumbs is not the only homophilic interactor that can exert such effects. Recent examples include E- and N-Cadherin patterning during eye morphogenesis in the fly (Chan et al., 2017), as well as tissue-specific expression of a Cadherin, Cad2, selectively in the neural cells in *Ciona robusta*, thereby patterning myosin activity at the neural/epidermal boundary where there is a step change in Cad2 expression (Hashimoto and Munro, 2018). Interestingly, in *Ciona* Cad2 is also titrated away from the tissue boundary due to homophilic interactions and also exerts a negative regulatory effect on myosin II accumulation. In this context though, and in contrast to the salivary gland placode boundary, it is RhoA activity that is polarised and increased at the tissue boundary, due to a selective recruitment of a RhoGAP by Cad2 (Hashimoto and Munro, 2018). Thus, although common principles in patterning tissue boundaries and patterning of cytoskeletal activity are repeatedly used in development, the fine molecular details vary depending on the tissue context.

We identified a mechanism that allows for a negative effect on Rok and myosin accumulation at membranes only when a pre-existing molecular anisotropy of an upstream factor is detected, thereby circumventing an on/off negative regulatory interaction. In the *Ciona* example above, Cad2 levels are high in constricting neural cells that internalise into the embryo, very reminiscent of the situation in the salivary gland placode. Thus, the effect of Cad2 on myosin is not absolute here either. We suspect that such a mechanism of modulation of membrane residence time via affecting the k_off_ might be widely employed during tissue morphogenesis, as it allows for fine-tuning of cytoskeletal activity being integrated with other essential cell biological functions of the upstream regulators. In the case of Crumbs, its crucial role in maintaining apical-basal polarity and thereby epithelial integrity can be combined with its planar tissue patterning role.

In summary, this example of planar patterning in a morphogenetic process illustrates that the study of such processes needs to take the dynamic behaviour of components into account. The current exciting advances in light microscopy, live imaging and image quantification will be of crucial help to facilitate and transform such analyses in live tissues during development.

## Acknowledgements

The authors would like to thank the following people; for reagents and fly stocks: Debbie Andrew, Jennifer Zallen, Tony Harris, Nicholas Harden, Barry Thompson, Norbert Perrimon.

Work in the Röper lab is supported by the Medical Research Council (file reference number U105178780).

## Author contribution

Conceptualisation, K.R. and C.S.; Methodology, K.R., C.S. J.B., T.S.; Software, J.B., T.S.; Investigation, K.R., C.S.; Resources, C.S., L.J.; Writing-Original Draft, K.R., C.S.; Funding Acquisition, K.R.

---

**Supplementary Figure 1, related to Figure 2.**
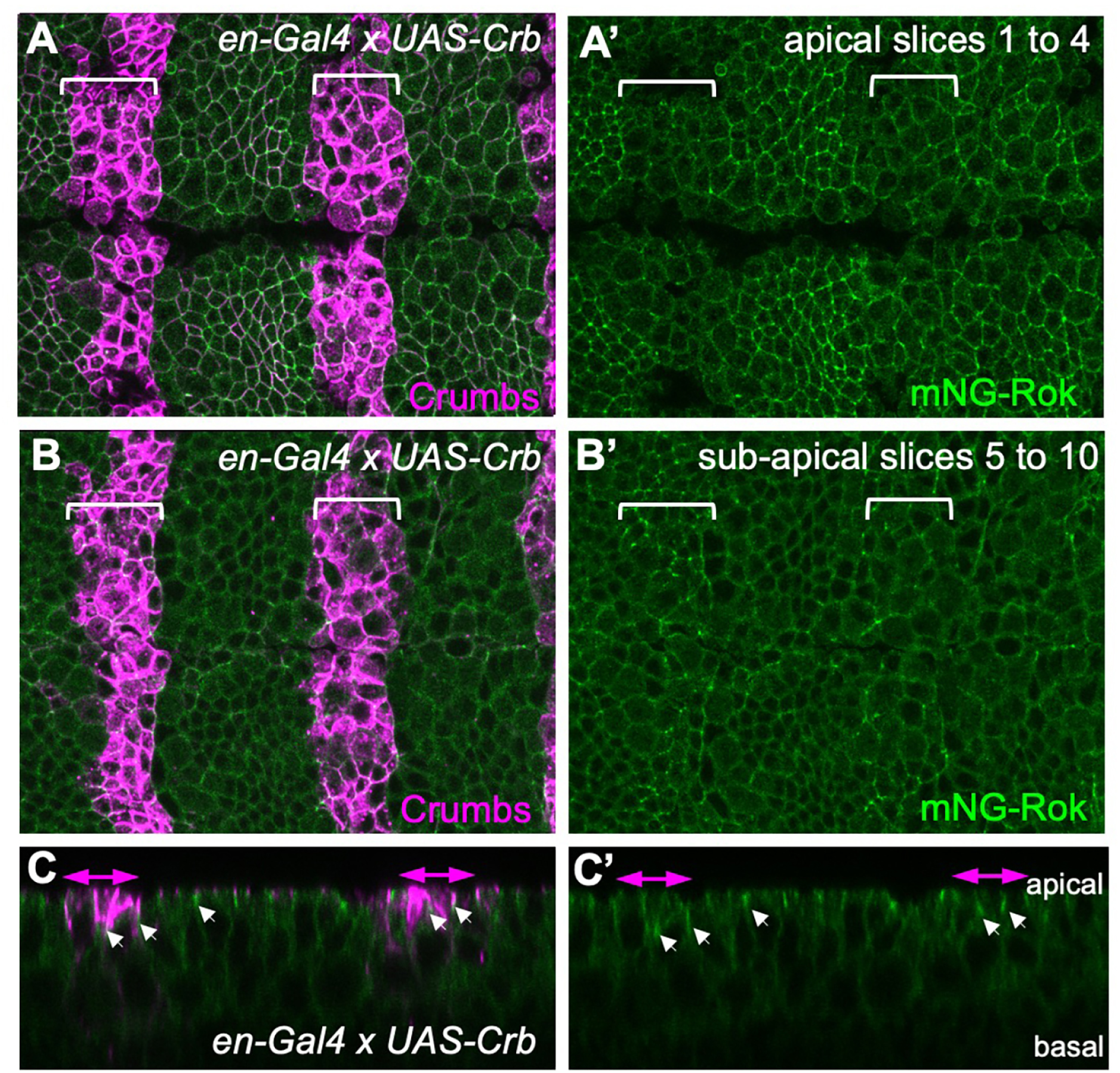
Crumbs membrane localisation locally affects Rok membrane localisation. Overexpression of Crumbs in stripes, using *en-Gal4 × UAS-Crumbs*, within the epidermis leads to spreading of Crumbs (magenta In **A-C**) to more basal position along the lateral plasma membrane. This in turn leads to a basal displacement of Rok (green) along the lateral side, visible through higher mNG-Rok membrane levels at a sub-apical position (**B’**) and loss of it at the apical-most lateral position (**A’**).

**Supplementary Figure 2, related to Figure 3.**
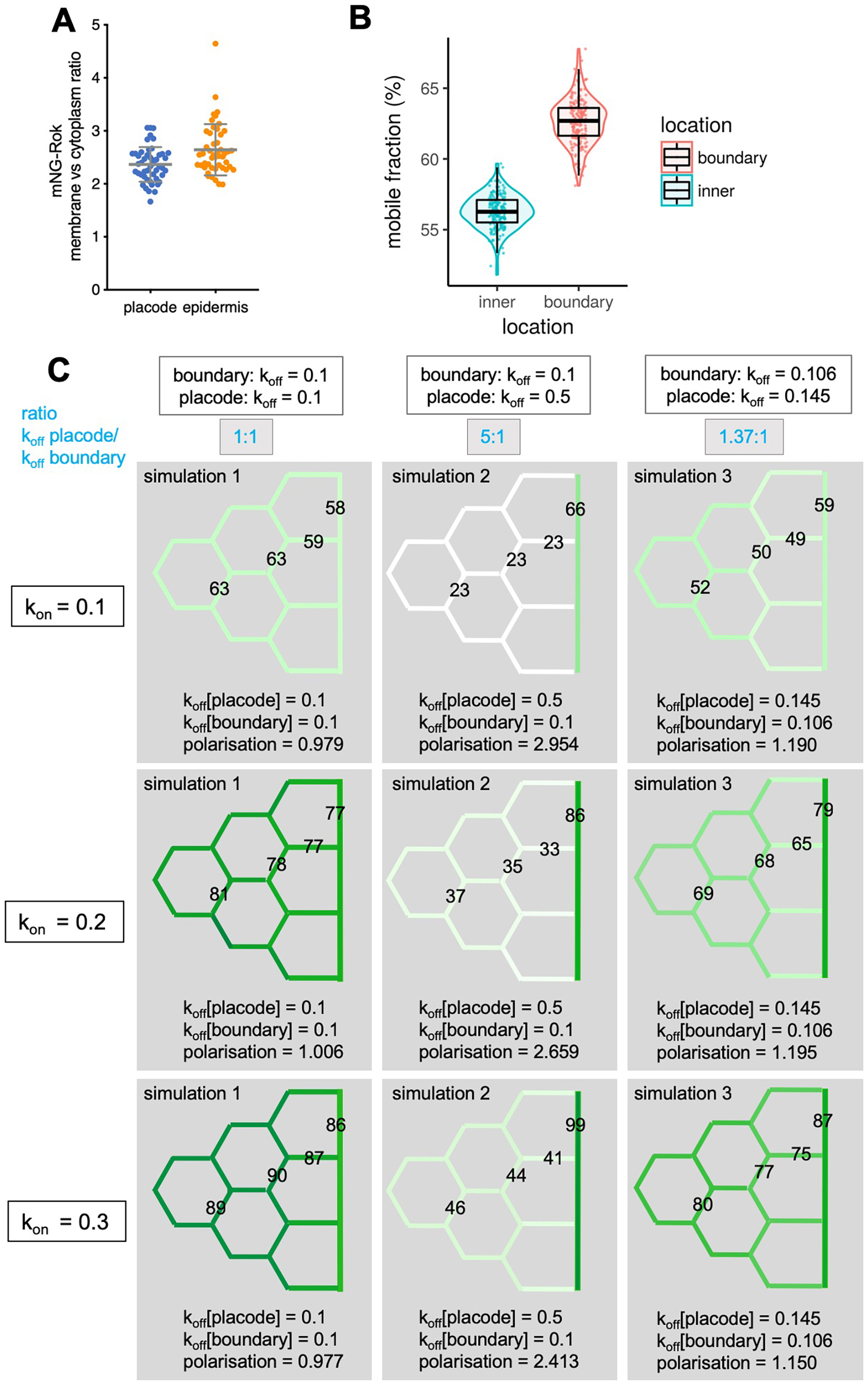
FRAP analysis and in silico simulation of Rok dynamics at the placode boundary versus inside the placode. **A** Membrane versus cytoplasm enrichment of mNG-Rok in inner placodal cells and surrounding epidermal cells. Data are represented as data points, mean and SEM. **B** Mobile fraction of mNG-Rok at placode inner membranes (blue; value is 0.56) or the boundary (red; value is 0.63), data points, mean and spread are shown. **C** Outputs of *in silico* simulations as in Fig. 3G, comparing the same combinations of k_off_ values paired with different k_on_ values (0.1; 0.2; 0.3). Color code for particle enrichment at junctions as in Fig. 3G, numbers on representative junctions are the particle numbers derived from simulations.

**Supplemental Figure 3, related to Figure 5.**
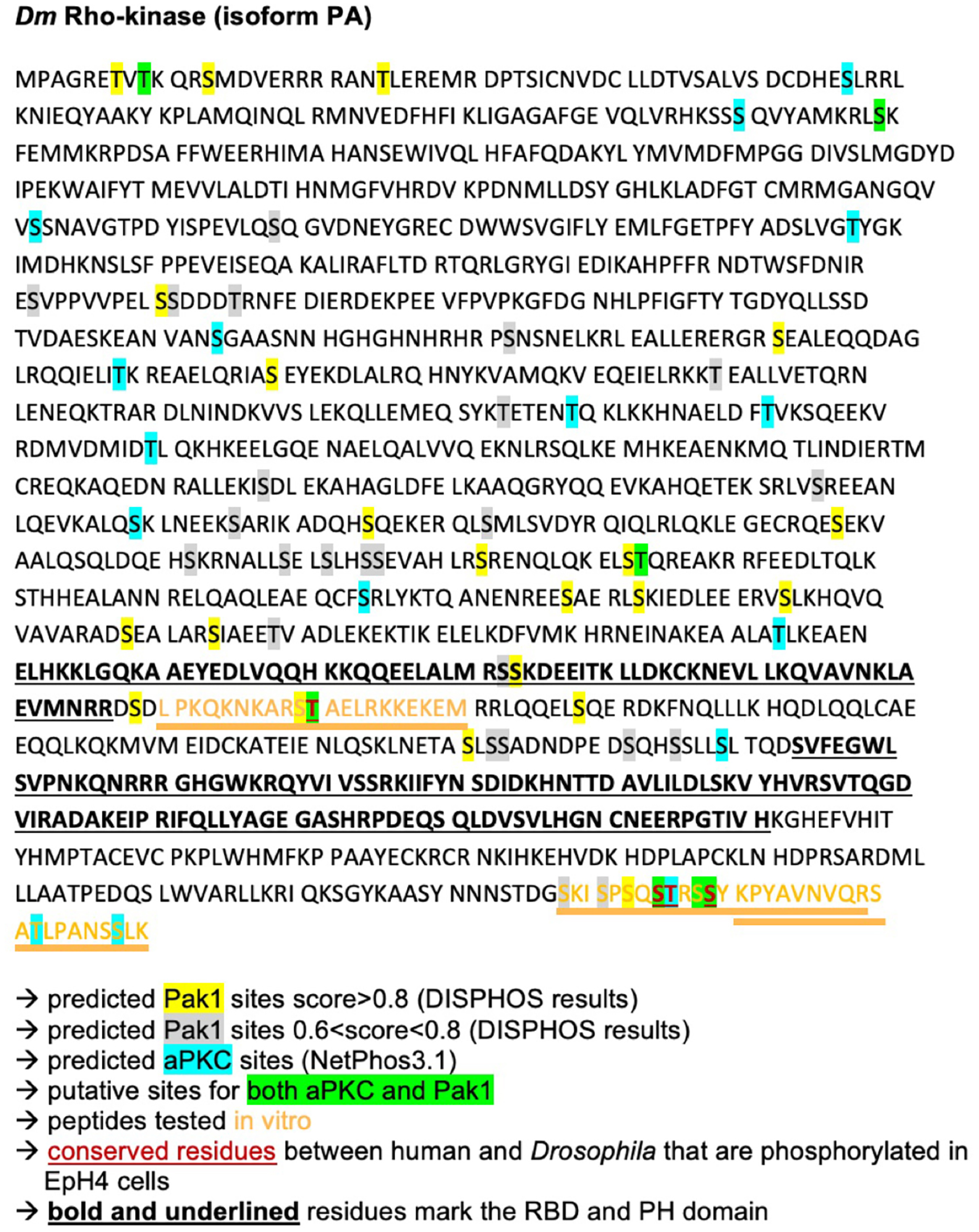
Analysis of predicted aPKC and Pak1 phosphorylation sites in Dm Rok. We used published predictive algorithms to identify potential aPKC and Pak1 phosphorylation sites in *Dm* Rok. Residues marked in yellow are predicted Pak1 sites (using DISPHOS) with a score >0.8, grey residues are predicted Pak1 sites (using DISPHOS) with a score between 0.6 and 0.8. Blue marks residues as predicted aPKC target sites (using NetPhos3.1). Green residues are predicted target sites for both aPKC and Pak1. Bold, underlined residues mark the RBD and PH domain. Conserved residues between human ROCK1 and *Dm* Rok that were found to be phosphorylated in EpH4 cells are shown in red and underlined (Ishiuchi and Takeichi, 2011).

**Supplemental Movie 1. Dynamics of Crumbs and Myosin at the salivary gland placode boundary.**

Time-lapse movie of an embryo with endogenously tagged Crumbs (green, Crumbs-GFP) and endogenously tagged Myosin II heavy chain (magenta, Zipper-YFP). Frames are 3 min apart.

**Supplemental Movie 2. Example of FRAP at a salivary gland placode boundary junction.**

Time-lapse movie of an embryo with endogenously tagged Rok (mNG-Rok), showing a bleach and recovery at the boundary of the placode, quantification in Figure 3.

**Supplemental Movie 3. Example of FRAP at a salivary gland inner cell junction.**

Time-lapse movie of an embryo with endogenously tagged Rok (mNG-Rok), showing a bleach and recovery at a junction of an inner placodal cell, quantification in Figure 3.

## Methods

### Fly stocks

The following transgenic fly lines were used: *sqhAX3; sqh::sqhGFP42* (Royou et al., 2004), *Daughterless-Gal4* and *enGal4* (Bloomington Stock Centre); *fkhGal4* (Henderson and Andrew, 2000; Zhou et al., 2001) [kin (Sivasankar et al., 2009)d gift of Debbie Andrew]; *Crb-GFP* (Huang et al., 2008); *Zip-YFP* (Lowe et al., 2014); *UAS-Crb* (Wodarz et al., 1995); *UAS-Venus-Rok[K116A], UAS-Venus-Rok[K116A]ΔRBD, UAS-Venus-Rok[K116A] ΔCC-SBD; UAS-Venus-Rok[K116A]ΔPH* (Simoes Sde et al., 2014); *Ubi-EGFP-AnillinRBD* (Munjal et al., 2015); *sfGFP-RhoGEF2* (Sarov et al., 2016); *UAS-aPKC[CAAX]* (Sotillos et al., 2004); *UAS-Pak1[myr]*; *UAS-Cdc42[V12]*; *sqh::Cdc42-mCherry* (Bloomington Stock Centre); *pak1[14]* (gift from B.Thompson); *y[1] sc[1] v[1]; [y[+t7.7] v[+t1.8]=nanos-Cas9]attp2* (gift from N. Perrimon).

### Generation of transgenic fly lines

To generate *Drosophila rok* transgenic lines, donor (150 ng/μl) and guide RNA (100 ng/μl) plasmids were injected in pools (Bischof et al., 2013) into *nanos::Cas9* (chromosome 3) embryos for the endogenous tagging, or into *mNG-Rok*; *nanos::Cas9* embryos for the generation of the phospho-site mutant.

#### Fluorescent tagging of endogenous Rok via homology directed repair using CRISPR-Cas9

Two gRNAs targeting loci near the start codon of the Rho kinase gene were cloned into pCFD3 vector (Addgene 49410) following the protocol from (Port et al., 2014). A step by step protocol is available at (www.crisprflydesign.org).

Sequences of the guide RNAs were as follows:

gRNA 1: GACCAACAGGAAGCAGCAGCTGG

gRNA2: GCGCCGGTGAGTGCACGAGATGG

PCR primers for cloning into pCFD3:

P56F: 5’-GTCGACCAACAGGAAGCAGCAGC-3’

P56R: 5’-AAACGCTGCTGCTTCCTGTTGGT-3’

P57F: 5’-GTCGCGCCGGTGAGTGCACGAGA-3’

P57R: 5’-AAACTCTCGTGCACTCACCGGCG-3’

A donor construct containing the mNeonGreen sequence in fusion with the *rok* gene in its genomic region was cloned into pBluescript SK(+) using Gibson assembly. *mNeonGreen* was cloned between two 1kb-long homology sequences corresponding to the genomic sequence on either side of the insertion site, to create homology arms for directed repair.The 1 kb regions were amplified by PCR on each side of the desired insertion site from genomic DNA. The mNeonGreen gene (Shaner et al., 2013), was amplified by PCR from the mNeonGreen vector (Allele Biotechnology), with the exclusion of the stop codon and the addition of a C-terminal linker.

Primers were designed with additional 5’ sequences (underlined below) to allow triple ligation of the three PCR products into a pBluescript SK(+) vector using the Gibson Assembly Master Mix (NEB). PAM sites were mutagenized (indicated in bold below) to prevent re-cutting by Cas9 after transgenesis.

Primer sequences:

Left homology arm PCR primers:

P51F: 5’<u>GACGGTATCGATAAGCTTGATATCG</u>GCGCAGCGTCTAATTGAAAC-3’

P51R: 5’GCTGATACTGCTGCT**aCA**GCTGCTGC-3’

Right homology arm PCR primers:

5’<u>TGCCAGCTGGACGAGAAACTGTGACCAAGC</u>AGCGCAGCATGGATGTGGAACGAAGGCGCCGgtgagtgcacgaga**tgt**cggcccaaaagc

mNeonGreen PCR primers:

P54F: 5’<u>GGAAGCAGCAGC**TGt**AGC</u>AGCAGTATCAGCTTGTTATCTTGCATTTGCATGGT GAGCAAGGGCGAGGAG-3’

P54R: 5’<u>GCTTGGTCACAGTTTCTCGTCCAGCTGGCA</u>TGCCGGATCCGCCGCCCGATCC GCCGCCGGATCCGCCCTTGTAAAGTTCATCCATCCCC-3’

Modifications were verified by sequencing of genomic DNA.

#### Mutagenesis of putative P-sites in endogenous mNG-rok via homology directed repair using CRISPR-Cas9

Guide RNAs:

Four gRNAs targeting loci on both sides of the C-terminal region of Rok containing the putative phosphorylation sites to be mutated were cloned into pCFD3 vector (Addgene 49410) following the protocol from (Port et al., 2014).

Sequences: LJ20, LJ21, LJ22, LJ23

Donor plasmid:

A donor construct containing, between two 1kb-long homology arms, a Rok C-term region mutagenized on 13 selected putative phosphorylation sites, was cloned into pBluescript SK(+). The Rok C-term region was amplified by PCR and cloned into pBluescript SK(+)with primers designed to mutagenise the selected 13 putative phosphorylation sites (Fig. 5B) and the 4 PAM sites.

Modifications were verified by sequencing of genomic DNA.

### Embryo Immunofluorescence Labelling, Confocal, and Time-lapse imaging

Embryos were collected on apple juice-agar plates and processed for immunofluorescence using standard procedures. Briefly, embryos were dechorionated in 50% bleach, fixed in 10% EM-grade formaldehyde, and stained with primary and secondary antibodies in PBT (PBS plus 0.5% bovine serum albumin and 0.3% Triton X-100). anti-Crumbs and anti-E-Cadherin antibodies were obtained from the Developmental Studies Hybridoma Bank at the University of Iowa (DSHB); anti-Pak1 (Harden et al., 1996); anti-aPKC (Santa Cruz); rhodamine-coupled phalloidin (Molecular Probes). Secondary antibodies used were Alexa Fluor 488/Fluor 549/Fluor 649 coupled (Molecular Probes) and Cy3 and Cy5 coupled (Jackson ImmunoResearch Laboratories). Samples were embedded in Vectashield (Vectorlabs).

Images of fixed samples were acquired on an Olympus FluoView 1200 or a Zeiss 780 Confocal Laser scanning system as z-stacks to cover the whole apical surface of cells in the placode. Z-stack projections were assembled in ImageJ or Imaris (Bitplane), 3D rendering was performed in Imaris.

For live time-lapse experiments embryos of the genotype *Crumbs-GFP Zipper-YFP* or *mNG-Rok* were dechorionated in 50% bleach and extensively rinsed in water. Embryos were manually aligned and attached to heptane-glue coated coverslips and mounted on custom-made metal slides; embryos were covered using halocarbon oil 27 (Sigma) and viability after imaging after 24h was controlled prior to further data analysis. Time-lapse sequences were imaged under a 40x/1.3NA oil objective on an inverted Zeiss 780 Laser scanning system. Z-stack projections to generate movies in Supplementary Material were assembled in ImageJ.

### Quantification of boundary polarisation and membrane versus cytoplasm enrichment

Fluorescence intensity was determined in ImageJ using projections covering the apical junctional region (as determined by Crumbs or E-Cadherin staining). Using Crumbs labeling of placodes, boundary cells showing clear Crumbs anisotropy were identified and quantified. 3- to 5-pixel-wide lines to cover the width of junctions (depending on the resolution of the image) were drawn at the boundary and at the sides of the boundary cells (see Fig. 4F). The intensity was divided by the area covered for each junction to determine the intensity/pixel. Values for the boundary junction were divided by the values of the side junctions to determine two polarisaiton values per boundary cell.

Membrane versus cytoplasm enrichment was determined in ImageJ using projections covering the apical junctional region (as determined by Crumbs or E-Cadherin staining). 3- to 5-pixel-wide lines to cover the width of junctions (depending on the resolution of the image) were drawn at a cell junction and a comparable line was drawn across the apical cytoplasm. The intensity of each line was divided by the area covered to determine the intensity/pixel. Values for the cell junction were divided by the values of the cytoplasm to determine the membrane versus cytoplasm enrichment.

N values for quantifications are as follows: Figure 4F’: *UAS-Rok[K116A]*: 4 placodes (3 embryos), 15 cells (29 polarisation values); *UAS-Rok[K116A]ΔCC-SBD*: 6 placodes (3 embryos), 18 cells (36 pol. values); *UAS-Rok[K116A]ΔRBD*: 5 placodes (3 embryos), 16 cells (32 pol. values); *UAS-Rok[K116A]ΔPH:* 6 placodes (3 embryos), 16 cells (32 pol. values)/ Figure 4I: *Ubi-EGFP-AniRBD:* 4 placodes (3 embryos), 15 cells (30 pol. values); *RhoGEF2 sfGFP:* 3 placodes (3 embryos), 17 cells (34 pol. values)/ Figure 5C”:*UAS-aPKC[CAAX]:* 3 embryos (6 overexpression stripes), 30 cells; *Control for aPKC stripe:*3 embryos (6 control stripes), 30 cells/ Figure 5D”: *UAS-Pak1[myr]*: 4 embryos (7 overexpression stripes), 35 cells; *Control for Pak1 stripe:* 4 embryos (7 control stripes), 35 cells/ Figure 6E: *mNG-Rok ctrl:* 4 placodes (2 embryos), 23 cells (46 pol. values); *mNG-Rok in pak1[14]:* 3 placodes (2 embryos), 15 cells (30 pol. values); *mNG-Rok fkhGal4 UAS-Pak1[myr]:* 4 placodes (2 embryos), 26 cells (32 pol. values)/ Figure 6J: *mCherry-Cdc42:* 4 placodes (3 embryos), 24 cells (48 pol. values); *Crumbs:* 3 placodes (2 embryos), 14 cells (28 pol. values)/ Figure 7G: *mNG-Rok ctrl:* 5 placodes (5 embryos), 23 cells (44 pol. values); *mNG-Rok[Pmut]*: 4 placodes (4 embryos), 18 cells (34 pol. measurements). Statistical significance in comparisons was determined using unpaired t-test. Plots show data points, mean and SEM. p-values are indicated as ** being <0.005, *** being <0.0005, ****<0.0001, ns being not significant.

### Embryo Viability Assay

Embryos of the genotype *mNG-Rok* control or *mNG-Rok[Pmut]* were treated as for live imaging and mounted in separate sets of 100 embryos per experiment and let to develop at 18°C. After 48 hours hatched larvae, unfertilised embryos and developed but dead embryos were counted.

### FRAP imaging and analysis

Movies of the apical region of the salivary placode epithelium in stage 11 mNG-Rok embryos were acquired on an Andor Revolution XD Spinning Disk confocal with a 100X Objective (image size 512×512 pixels), 488 laser. 1.4 μm thick Z sections (5×0.35 μm) were acquired to compensate for movement in z during acquisition. For the FRAP, bleach dwell time was 500μs with 100% 488 laser power. Images were acquired at 1 second intervals, 6 time points pre-bleach and 40 time points post bleach.

Movies were analysed in ImageJ. Fluorescence intensity was measured in an 8-pixel circular ROI at the site of bleach.

Measures were then normalised to account for the general photobleaching caused by image acquisition. All values were multiplied by a photobleaching correction factor *C_photobleaching_*:

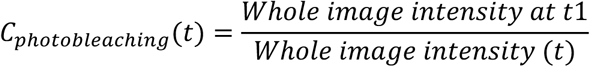

Normalised Fluorescence intensity measurements were used to plot the percentage of fluorescence recovery after photobleaching as follows: with *F*_*prebleach*_ = *avg F*_*t*1 *to t*6_

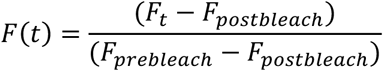

### FRAP curve fitting and statistical analysis

The k_off_ was estimated from the whole set of normalized fluorescence recovery curves. We modeled the recovery using a single exponential function in the form of: *a* (1 − *e*^−*k*_*off*_*t*^). A non-linear regression algorithm was used to estimate parameters from the data without prior averaging. As individual fluorescence recovery curves were noisy, we used a bootstrap procedure generating 200 estimates to estimate the statistical confidence of the estimated k_off_. This allowed us to compute a p-value using a t-test based on bootstrap variance. Note that this p-value does not depend on the number of bootstrap samples.

### In silico Rok particle simulation

As illustrated in Figure 3F, we simulated the diffusion of membrane bound and unbound particles (corresponding to Rok) within a pixel representation of a placodal cell layer containing inner and boundary cells. Our intention was to create a deliberately simple two-dimensional model that might nonetheless recapitulate the salient features of the observed planar polarity. Different membrane association and dissociation constants were modelled in different regions of the simulation according to the zones illustrated in Figure 3G, i.e. distinguishing the placode boundary from the inner placode membranes. Here the representation of the cell membranes with superimposed particle positions automatically provided a visualization of the simulation progress. By counting bound and unbound particles in different regions, polarisation at the boundary was quantified and compared with microscopic measurements. An exhaustive grid-search for parameters, which was initially corse and then more fine-grained, was performed to determine the combinations of diffusion, binding, unbinding and simulation time-step values that matched observations.

In detail, point particles were modelled as moving with random sequential displacements according to 2D Gaussian diffusion within a grid-based (i.e. pixel) representation of a cell layer with membrane boundaries. Particles were set to freely diffuse (off-grid) in the cell interior or diffuse laterally along the interior edge of the cell membrane; i.e. as unbound and bound sates. The positional variance for free diffusion (sigma) was set to correspond approximately to a diffusion coefficient of 25 μm^2^/s within a 25 μm wide cell area, which is the diffusion coefficient measured *in vivo* for mammalian GFP-ROCK2 (Truebestein et al., 2015). Diffusion of bound particles within the membrane was restricted to adjacent sites and was much slower than for free particles, with diffusion coefficients tested in the range 0.1 - 0.001 μm^2^/s (which made no practical difference) and final simulations set at 0.01 μm^2^/s. Using a probabilistic model, that can be related to k_on_ (association) and k_off_ (dissociation) constants, free particles that collided with the cell membrane were able to bind and bound particles were able to spontaneously unbind/dissociate. Here simulation steps typically corresponded to time segments of 50 ms, though a range of values was tested. k_off_ (in units of per second) was used to set the long-term probability of each bound particle spontaneously unbinding within the simulation time-step. k_on_ was less straightforward to model as, for particles which collide with the membrane, it depends on the concentration of receptive membrane sites and this varies throughout the simulation. Accordingly, at-membrane binding probabilities of initially free particles were set dynamically so that the derived, average k_on_ measured in the simulation converged to a desired value. In essence membrane binding probability was increased when the target k_on_ was undershot and reduced when overshot, averaging around a fixed k_on_. A single k_on_ value was used for all membrane regions, but different k_off_ values were used for placode boundary membranes and inner placode membranes.

The *in vivo* k_on_ used in simulations shown in Fig. 3 was estimated using the following equation from the “diffusion plus binding” model from (Sprague et al., 2004):

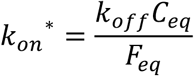

The k_off_ was calculated from our FRAP experiments, and we found that the 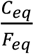 ratio could be deduced from parameters measured in vivo:

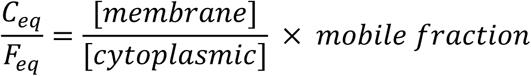

We measured the membrane to cytoplasmic ratio of mNG-Rok in stage 11 embryos and the value of the k_off_ and mobile fraction were calculated from fitted FRAP curves (Fig. 3C and Supplemental Fig.2 A,B): *k*_*off*_ = 0.14; mobile fraction = 0.56; membrane versus cytoplasm ration: 2.367.

Using the above equation and *in vivo* measured values, we estimated the k_on_ to be about 0.186, using 0.2 for simulations in Fig. 3, but also testing a range of k_on_ values between 0.1 and 0.3 in the simulations. Varying k_on_ within this range did not affect the polarisation values.

In order that the membrane binding sites be capable of saturation, the cell edge pixels were subjected to a maximum occupancy value. For the final simulations this was set at a value of 1 particle per pixel. Occupancy limits of 1-3 particles per pixel were tested and overall this made little difference to the cell polarity, but had a notable effect on the bound/free ratio, as we might expect.

Although the number of particles modelled within each cell could be varied, this made little difference to the long term bound/unbound ratios, but naturally more particles gave smoother, less variant values. Typically 1000 particles per cell were simulated. The particle simulation was started with randomly distributed particle positions within the cell interiors and progressed through 2000 unmonitored steps to equilibrate the model. Thereafter analyses of the particle positions were made at regularly spaced intervals for a further 10,000 steps. At each sample point the counts of bound and unbound particles were recorded for the regions marked in Figure 3G and later averaged for the whole simulation.

Python code to perform the 2D cell particle simulations, generating both regional counts and pixmap images, is available at: https://github.com/tjs23/memodis

### Rho-kinase sequence analysis

We used published predictive algorithms to identify potential aPKC and Pak1 phosphorylation sites in *Dm* Rok, DISPHOS (http://www.dabi.temple.edu/disphos/) and NetPhos3.1 (http://www.cbs.dtu.dk/services/NetPhos/). Rok sequences form different species were compared and aligned in EMBOSS Matcher (https://www.ebi.ac.uk/Tools/psa/emboss_matcher/).

### In vitro kinase assay

The following High-pressure liquid chromatography (HPLC)-purified peptides were ordered from Biomatik:

BazpepS980: EHFSRDALGRR**S**ISEKHHAAL

Rokpep1T: LPKQKNKARS**T**AELRKKEKEM

Rokpep2STS: SKISPSQ**ST**RS**S**YKPYAVNV

Rokpep2AAA: SKISPSQ**AA**RS**A**YKPYAVNV

Rokpep3S: KPYAVNVQRSATLPANS**S**LK

For in vitro kinase assays, 10μg of peptide substrate were incubated with either 150pg recombinant human Pak1 kinase domain (AbNOVA) or 0.1 μM recombinant human PKCι kinase domain (a gift from N. McDonald via B. Thompson) for 30 min at 30°C in kinase reaction buffer (50 mM HEPES [(4-(2-hydroxyethyl)-1-piperazineethanesulfonic acid] pH 7.5, 10 mM MgCl2, 1 mM EGTA, 0.01% Brij35) containing 10 μM cold ATP and 3 μCi γ-P33 ATP (Hartmann Analytic GmbH). Samples were blotted on 2cm × 2cm squares of P81 phosphocellulose paper (Millipore) and washed 3 × 10 min in 1% phosphoric acid, then 5 min in acetone. Dried papers were then transferred to scintillation vials and immersed in liquid scintillation cocktail (Ultima Gold XR, Perkin Elmer). Incorporation of γ-P33 was quantified in counts per minute by scintillation counting (Beckman LS 6500).

## References

Aguilar-Aragon, M., Elbediwy, A., Foglizzo, V., Fletcher, G.C., Li, V.S.W., and Thompson, B.J. (2018). Pak1 Kinase Maintains Apical Membrane Identity in Epithelia. Cell Reports 22, 1639–1646.

Amano, M., Chihara, K., Nakamura, N., Kaneko, T., Matsuura, Y., and Kaibuchi, K. (1999). The COOH terminus of Rho-kinase negatively regulates rho-kinase activity. J. Biol. Chem. 274, 32418–32424.

Amano, M., Nakayama, M., and Kaibuchi, K. (2010). Rho-kinase/ROCK: A key regulator of the cytoskeleton and cell polarity. Cytoskeleton 67, 545–554.

Bahri, S., Wang, S., Conder, R., Choy, J., Vlachos, S., Dong, K., Merino, C., Sigrist, S., Molnar, C., Yang, X., et al. (2010). The leading edge during dorsal closure as a model for epithelial plasticity: Pak is required for recruitment of the Scribble complex and septate junction formation. Development 137, 2023–2032.

Bailey, M.J., and Prehoda, K.E. (2015). Establishment of Par-Polarized Cortical Domains via Phosphoregulated Membrane Motifs. Dev. Cell 35, 199–210.

Bischof, J., Bjorklund, M., Furger, E., Schertel, C., Taipale, J., and Basler, K. (2013). A versatile platform for creating a comprehensive UAS-ORFeome library in Drosophila. Development 140, 2434–2442.

Blom, N., Gammeltoft, S., and Brunak, S. (1999). Sequence and structure-based prediction of eukaryotic protein phosphorylation sites. J. Mol. Biol. 294, 1351–1362.

Blom, N., Sicheritz-Ponten, T., Gupta, R., Gammeltoft, S., and Brunak, S. (2004). Prediction of post-translational glycosylation and phosphorylation of proteins from the amino acid sequence. Proteomics 4, 1633–1649.

Bokoch, G.M. (2003). Biology of the p21-activated kinases. Annual Review of Biochemistry 72, 743–781.

Booth, A.J., Blanchard, G.B., Adams, R.J., and Röper, K. (2014). A dynamic microtubule cytoskeleton directs medial actomyosin function during tube formation. Dev. Cell 29, 562–576.

Brand, A.H., and Perrimon, N. (1993). Targeted gene expression as a means of altering cell fates and generating dominant phenotypes. Development 118, 401–415.

Bulgakova, N.A., and Knust, E. (2009). The Crumbs complex: from epithelial-cell polarity to retinal degeneration. J. Cell Sci. 122, 2587–2596.

Calzolari, S., Terriente, J., and Pujades, C. (2014). Cell segregation in the vertebrate hindbrain relies on actomyosin cables located at the interhombomeric boundaries. EMBO J. 33, 686–701.

Castelli-Gair Hombria, J. (2016). Organogenetic Gene Networks (Springer).

Chan, E.H., Chavadimane Shivakumar, P., Clement, R., Laugier, E., and Lenne, P.F. (2017). Patterned cortical tension mediated by N-cadherin controls cell geometric order in the Drosophila eye. eLife 6.

Chang, L.H., Chen, P., Lien, M.T., Ho, Y.H., Lin, C.M., Pan, Y.T., Wei, S.Y., and Hsu, J.C. (2011). Differential adhesion and actomyosin cable collaborate to drive Echinoid-mediated cell sorting. Development 138, 3803–3812.

Conder, R., Yu, H., Zahedi, B., and Harden, N. (2007). The serine/threonine kinase dPak is required for polarized assembly of F-actin bundles and apical-basal polarity in the Drosophila follicular epithelium. Dev. Biol. 305, 470–482.

Dahmann, C., and Basler, K. (1999). Compartment boundaries: at the edge of development. Trends Genet. 15, 320–326.

Fletcher, G.C., Lucas, E.P., Brain, R., Tournier, A., and Thompson, B.J. (2012). Positive feedback and mutual antagonism combine to polarize crumbs in the Drosophila follicle cell epithelium. Curr. Biol. 22, 1116–1122.

Galea, G.L., Cho, Y.J., Galea, G., Mole, M.A., Rolo, A., Savery, D., Moulding, D., Culshaw, L.H., Nikolopoulou, E., Greene, N.D.E., et al. (2017). Biomechanical coupling facilitates spinal neural tube closure in mouse embryos. Proc. Natl. Acad. Sci. USA 114, E5177–E5186.

Girdler, G.C., and Röper, K. (2014). Controlling cell shape changes during salivary gland tube formation in Drosophila. Semin. Cell Dev. Biol. 31, 74–81.

Harden, N., Lee, J., Loh, H.Y., Ong, Y.M., Tan, I., Leung, T., Manser, E., and Lim, L. (1996). A Drosophila homolog of the Rac- and Cdc42-activated serine/threonine kinase PAK is a potential focal adhesion and focal complex protein that colocalizes with dynamic actin structures. Mol. Cell Biol. 16, 1896–1908.

Hashimoto, H., and Munro, E. (2018). Differential expression and homotypic enrichment of a classic Cadherin directs tissue-level contractile asymmetry during neural tube closure. BioRxiv.

Henderson, K.D., and Andrew, D.J. (2000). Regulation and function of Scr, exd, and hth in the Drosophila salivary gland. Dev. Biol. 217, 362–374.

Hong, Y. (2018). aPKC: the Kinase that Phosphorylates Cell Polarity. F1000Res 7.

Huang, J., Zhou, W., Watson, A.M., Jan, Y.N., and Hong, Y. (2008). Efficient ends-out gene targeting in Drosophila. Genetics 180, 703–707.

Ishiuchi, T., and Takeichi, M. (2011). Willin and Par3 cooperatively regulate epithelial apical constriction through aPKC-mediated ROCK phosphorylation. Nat. Cell Biol. 13, 860–866.

Jacinto, A., Wood, W., Woolner, S., Hiley, C., Turner, L., Wilson, C., Martinez-Arias, A., and Martin, P. (2002). Dynamic analysis of actin cable function during Drosophila dorsal closure. Curr. Biol. 12, 1245–1250.

Julian, L., and Olson, M.F. (2014). Rho-associated coiled-coil containing kinases (ROCK): structure, regulation, and functions. Small GTPases 5, e29846.

Lowe, N., Rees, J.S., Roote, J., Ryder, E., Armean, I.M., Johnson, G., Drummond, E., Spriggs, H., Drummond, J., Magbanua, J.P., et al. (2014). Analysis of the expression patterns, subcellular localisations and interaction partners of Drosophila proteins using a pigP protein trap library. Development 141, 3994–4005.

Major, R.J., and Irvine, K.D. (2005). Influence of Notch on dorsoventral compartmentalization and actin organization in the Drosophila wing. Development 132, 3823–3833.

Major, R.J., and Irvine, K.D. (2006). Localization and requirement for Myosin II at the dorsal-ventral compartment boundary of the Drosophila wing. Dev. Dyn. 235, 3051–3058.

Mason, F.M., Xie, S., Vasquez, C.G., Tworoger, M., and Martin, A.C. (2016). RhoA GTPase inhibition organizes contraction during epithelial morphogenesis. J. Cell Biol. 214, 603–617.

mod, E.C., Roy, S., Ernst, J., Kharchenko, P.V., Kheradpour, P., Negre, N., Eaton, M.L., Landolin, J.M., Bristow, C.A., Ma, L., et al. (2010). Identification of functional elements and regulatory circuits by Drosophila modENCODE. Science 330, 1787–1797.

Monier, B., Pelissier-Monier, A., Brand, A.H., and Sanson, B. (2010). An actomyosin-based barrier inhibits cell mixing at compartmental boundaries in Drosophila embryos. Nat. Cell Biol. 12, 60–65; sup pp 61-69.

Munjal, A., Philippe, J.M., Munro, E., and Lecuit, T. (2015). A self-organized biomechanical network drives shape changes during tissue morphogenesis. Nature 524, 351–355.

Nakamura, M., Verboon, J.M., and Parkhurst, S.M. (2017). Prepatterning by RhoGEFs governs Rho GTPase spatiotemporal dynamics during wound repair. J. Cell Biol. 216, 3959–3969.

Pare, A.C., Vichas, A., Fincher, C.T., Mirman, Z., Farrell, D.L., Mainieri, A., and Zallen, J.A. (2014). A positional Toll receptor code directs convergent extension in Drosophila. Nature 515, 523–527.

Pirraglia, C., Walters, J., and Myat, M.M. (2010). Pak1 control of E-cadherin endocytosis regulates salivary gland lumen size and shape. Development 137, 4177–4189.

Port, F., Chen, H.M., Lee, T., and Bullock, S.L. (2014). Optimized CRISPR/Cas tools for efficient germline and somatic genome engineering in Drosophila. Proc. Natl. Acad. Sci. USA 111, E2967–2976.

Ramkumar, N., Omelchenko, T., Silva-Gagliardi, N.F., McGlade, C.J., Wijnholds, J., and Anderson, K.V. (2016). Crumbs2 promotes cell ingression during the epithelial-to-mesenchymal transition at gastrulation. Nat. Cell Biol. 18, 1281–1291.

Rane, C.K., and Minden, A. (2014). P21 activated kinases: structure, regulation, and functions. Small GTPases 5.

Rennefahrt, U.E., Deacon, S.W., Parker, S.A., Devarajan, K., Beeser, A., Chernoff, J., Knapp, S., Turk, B.E., and Peterson, J.R. (2007). Specificity profiling of Pak kinases allows identification of novel phosphorylation sites. J. Biol. Chem. 282, 15667–15678.

Röper, K. (2012). Anisotropy of Crumbs and aPKC Drives Myosin Cable Assembly during Tube Formation. Dev. Cell 23, 939–953.

Röper, K. (2013). Supracellular actomyosin assemblies during development. Bioarchitecture 3, 45–49.

Royou, A., Field, C., Sisson, J.C., Sullivan, W., and Karess, R. (2004). Reassessing the role and dynamics of nonmuscle myosin II during furrow formation in early Drosophila embryos. Mol. Biol. Cell 15, 838–850.

Sanchez-Corrales, Y.E., Blanchard, G.B., and Roper, K. (2018). Radially patterned cell behaviours during tube budding from an epithelium. eLife 7. http://doi.org/10.7554/eLife.35717

Sarov, M., Barz, C., Jambor, H., Hein, M.Y., Schmied, C., Suchold, D., Stender, B., Janosch, S., K, J.V., Krishnan, R.T., et al. (2016). A genome-wide resource for the analysis of protein localisation in Drosophila. eLife 5, e12068.

Shaner, N.C., Lambert, G.G., Chammas, A., Ni, Y., Cranfill, P.J., Baird, M.A., Sell, B.R., Allen, J.R., Day, R.N., Israelsson, M., et al. (2013). A bright monomeric green fluorescent protein derived from Branchiostoma lanceolatum. Nat. Methods 10, 407–409.

Sidor, C., and Röper, K. (2016). Genetic Control of Salivary Gland Tubulogenesis in Drosophila. In Organogenetic Gene Networks, (J. Castelli-Gair Hombría, and P. Bovolenta, eds.) Springer International Publishing, pp. 125–149.

Simoes Sde, M., Blankenship, J.T., Weitz, O., Farrell, D.L., Tamada, M., Fernandez-Gonzalez, R., and Zallen, J.A. (2010). Rho-kinase directs Bazooka/Par-3 planar polarity during Drosophila axis elongation. Dev. Cell 19, 377–388.

Simoes Sde, M., Mainieri, A., and Zallen, J.A. (2014). Rho GTPase and Shroom direct planar polarized actomyosin contractility during convergent extension. J. Cell Biol. 204, 575–589.

Sivasankar, S., Zhang, Y., Nelson, W.J., and Chu, S. (2009). Characterizing the initial encounter complex in cadherin adhesion. Structure 17, 1075–1081.

Sotillos, S., Diaz-Meco, M.T., Caminero, E., Moscat, J., and Campuzano, S. (2004). DaPKC-dependent phosphorylation of Crumbs is required for epithelial cell polarity in Drosophila. J. Cell Biol. 166, 549–557.

Sprague, B.L., Pego, R.L., Stavreva, D.A., and McNally, J.G. (2004). Analysis of binding reactions by fluorescence recovery after photobleaching. Biophys. J. 86, 3473–3495.

Tepass, U. (2012). The apical polarity protein network in Drosophila epithelial cells: regulation of polarity, junctions, morphogenesis, cell growth, and survival. Annu. Rev. Cell Dev. Biol. 28, 655–685.

Tepass, U., Godt, D., and Winklbauer, R. (2002). Cell sorting in animal development: signalling and adhesive mechanisms in the formation of tissue boundaries. Curr. Opin. Genet. Dev. 12, 572–582.

Tetley, R.J., Blanchard, G.B., Fletcher, A.G., Adams, R.J., and Sanson, B. (2016). Unipolar distributions of junctional Myosin II identify cell stripe boundaries that drive cell intercalation throughout Drosophila axis extension. eLife 5.

Truebestein, L., Elsner, D.J., Fuchs, E., and Leonard, T.A. (2015). A molecular ruler regulates cytoskeletal remodelling by the Rho kinases. Nature Communications 6, 10029.

Truebestein, L., Elsner, D.J., and Leonard, T.A. (2016). Made to measure - keeping Rho kinase at a distance. Small GTPases 7, 82–92.

Welch, H., Eguinoa, A., Stephens, L.R., and Hawkins, P.T. (1998). Protein kinase B and rac are activated in parallel within a phosphatidylinositide 3OH-kinase-controlled signaling pathway. J. Biol. Chem. 273, 11248–11256.

Wodarz, A., Hinz, U., Engelbert, M., and Knust, E. (1995). Expression of crumbs confers apical character on plasma membrane domains of ectodermal epithelia of Drosophila. Cell 82, 67–76.

Zhou, B., Bagri, A., and Beckendorf, S.K. (2001). Salivary gland determination in Drosophila: a salivary-specific, fork head enhancer integrates spatial pattern and allows fork head autoregulation. Dev. Biol. 237, 54–67.

Zou, J., Wang, X., and Wei, X. (2012). Crb Apical Polarity Proteins Maintain Zebrafish Retinal Cone Mosaics via Intercellular Binding of Their Extracellular Domains. Dev. Cell 22, 1261–1274.

